# CoCUT&Tag maps linked chromatin states at single-molecule, single-cell resolution

**DOI:** 10.64898/2026.04.27.721191

**Authors:** Wenjie Sun, Emma Baird, Harry J. Ashbaugh, Alexandria P. Eiken, Derek H. Janssens

## Abstract

Genomic methods for profiling multiple chromatin features face tradeoffs in specificity, sensitivity, and the ability to measure chromatin co-occupancy in rare cell types and primary tissues. Here, we present CoCUT&Tag, which uses synthetic antigen-peptide repeats paired with nanobodies to enhance Tn5 transposase targeting and enable single-cell measurement of the co-occupancy of two chromatin features on the same DNA molecule. We apply CoCUT&Tag to test the model that bivalent chromatin, marked by the coexistence of H3K27me3 and H3K4 methylation, preserves stem-cell multipotency by keeping developmental genes poised for future activation. We find that during human hematopoietic differentiation, global bivalency increases, whereas the number of genes in an active chromatin state is highest in stem and multipotent progenitor cells. Bivalent and unmarked genes activate with similar timing, but bivalent genes define a distinct regulatory class that achieves higher expression, is associated with more numerous and stronger enhancers, and is preferentially derepressed following loss of Polycomb-repressor proteins. Our study shows that bivalent chromatin increases enhancer demand to enable high-amplitude gene activation, demonstrating that CoCUT&Tag can resolve longstanding questions about combinatorial chromatin regulation.

## Introduction

Gene regulation is controlled by a complex milieu of nuclear proteins that function together to regulate transcription from the same DNA template. However, due to technical challenges, most genomic assays continue to profile one chromatin protein at a time. A fundamental limitation of this approach is that overlapping genomic binding sites for two proteins do not necessarily mean that they occupy the same sites in the same cells, much less at the same time. This makes direct measurement of linked chromatin states essential for understanding how combinations of chromatin features encode regulatory function.

Even genomic assays that profile two chromatin features at a time remain limited. Bulk approaches such as sequential ChIP and Co-ChIP can detect co-localized histone marks genome-wide^1,2^. However, in heterogeneous tissues, these approaches cannot determine whether linked chromatin states observed at different loci arise from the same cells. Single-cell “multifactorial” methods for joint profiling of two chromatin proteins face a different set of tradeoffs between the specificity, sensitivity, and direct measurement of co-occupancy. Some rely on shared secondary reagents or protein A-based tethering strategies that risk cross-reaction between targets^3-5^. Other methods use antibody-specific Tn5 fusions to preserve direct tethering but yield relatively sparse signal^6^, or improve read recovery at the cost of losing direct information about whether two features occupy the same DNA molecules^7,8^. As a result, combinatorial chromatin states in rare cell types and primary tissues are still often inferred from overlapping profiles rather than measured directly.

To overcome these limitations, we developed co-occupancy CUT&Tag (CoCUT&Tag), a chromatin profiling strategy that uses orthogonal antigen-peptide repeats and nanobody fusions to amplify signal from two chromatin targets while preserving their linkage on DNA. In CoCUT&Tag, rabbit and mouse primary antibodies direct orthogonal repeated-antigen proteins to two chromatin features, where they act as scaffolds to increase the number of barcoded Tn5 transposases tethered at each site through fused antigen-recognition nanobodies. This design unambiguously resolves target-specific binding sites and regions co-occupied by two chromatin features on the same DNA molecule. In addition, CoCUT&Tag is compatible with automated single-cell combinatorial-indexing workflows and generates high-quality linked chromatin profiles from 25,000-50,000 cells per run of a nanowell dispenser.

We applied CoCUT&Tag to investigate the longstanding model that genes controlling cell-fate decisions are held in a bivalent chromatin state that keeps them silent but poised for future activation^9,10^. Bivalent chromatin is defined by the coexistence of the Polycomb-associated repressive mark H3K27me3 and the Trithorax-associated activating mark H3K4me3^1^, and has been particularly influential in stem cell biology, where it is thought to preserve the multipotency of stem cells to generate daughter cells that differentiate to acquire one of multiple distinct cell fates. Despite extensive investigation, the functional significance of bivalent chromatin remains unresolved. For example, during differentiation, bivalent genes are not necessarily activated earlier than genes that begin in unmarked or repressed states^11^, raising the possibility that bivalency does not simply poise genes for activation, and may serve a regulatory function that has yet to be defined.

Recent perturbation studies further highlight the functional importance of bivalent chromatin. During hematopoiesis, loss of H3K4 methylation causes severe differentiation defects across multiple lineages, and simultaneously reducing H3K27 methylation partially rescues these phenotypes^12^. These findings indicate that bivalent chromatin reflects a functional balance between activating and repressive inputs. However, it remains unclear why some developmental genes retain both inputs simultaneously, rather than remaining unmarked until activation is required. This raises the possibility that bivalency is more than a transient intermediate between active and repressed chromatin, and instead reflects a stable identity encoded at specific developmental genes.

Despite the association with stem-cell multipotency, bivalent chromatin is not confined to stem cell populations and is also found in differentiated cell types and adult tissues^13,14^, where Polycomb is required to maintain repression of bivalent loci^15^. Together, these observations suggest bivalency acts as a more general regulatory state, with important functions beyond the context of stem-cell multipotency.

We find that global bivalency increases during human hematopoietic differentiation and that bivalent genes are not activated earlier than other genes, but define a distinct regulatory class associated with stronger enhancer input and higher levels of gene expression once activated. We further identify bivalent genes that remain transcriptionally silent despite nearby lineage-restricted H3K4me1-marked enhancer-like features. These loci are preferentially derepressed following loss of the Polycomb-group protein BMI1, indicating that Polycomb increases the enhancer input required for gene expression. Together, these findings support an enhancer-threshold model in which Polycomb-mediated repression restrains bivalent genes until sufficient regulatory input accumulates to drive high-amplitude expression.

## Results

### CoCUT&Tag identifies genomic regions bound by two proteins simultaneously

To directly measure co-occupancy of two chromatin features on the same DNA fragments, we developed CoCUT&Tag, a multifactorial CUT&Tag strategy built on two orthogonal repeated-antigen scaffolds that recruit Tn5 transposases loaded with distinct barcoded adapters (**Fig. 1A**). The SunTag and MoonTag are repeated antigen peptides recognized by specific nanobodies that were originally developed for signal amplification in live-cell imaging assays^16,17^. To adapt the SunTag and MoonTag for CoCUT&Tag, we fused them to anti-rabbit and anti-mouse IgG nanobodies^18^, respectively. In this configuration, rabbit and mouse primary antibodies independently target the SunTag and MoonTag scaffold proteins to distinct chromatin features, where they recruit anti-SunTag-Tn5 and anti-MoonTag-Tn5 fusion proteins (**Fig. 1A**). In the resulting libraries, single-barcode fragments report genomic regions bound by individual chromatin features, whereas mixed-barcode fragments identify genomic regions occupied by both chromatin features simultaneously (**Fig. 1A**). We engineered and purified the full CoCUT&Tag reagent set and optimized bulk reaction conditions for library quality and mappability (**Supplementary Fig. 1**).

**Figure 1.**
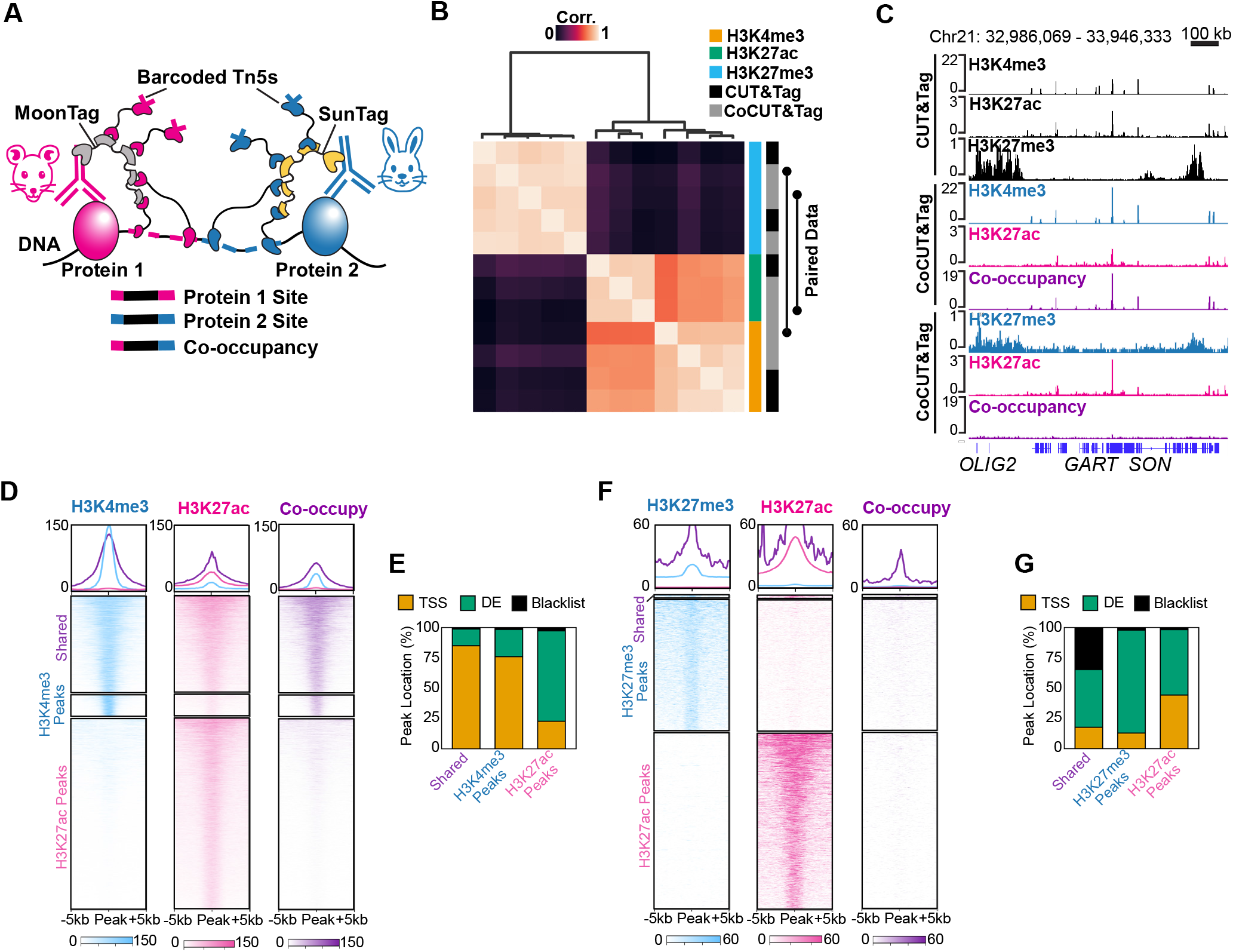
CoCUT&Tag identifies genomic regions bound by two proteins simultaneously. **(A)** Schematic of CoCUT&Tag. Two chromatin targets are recognized by rabbit- and mouse-derived primary antibodies, followed by orthogonal SunTag and MoonTag scaffolds. Target-specific Tn5 adapters generate single-barcode fragments, whereas mixed-barcode fragments recovered from the same locus report co-occupancy of the two chromatin features on the same DNA molecule. **(B)** Genome-wide correlation matrix comparing standard CUT&Tag and CoCUT&Tag. Right bars indicate CoCUT&Tag paired samples. **(C)** Genome browser tracks showing the specificity of CoCUT&Tag. Co-occupancy signal is detected at loci marked by H3K4me3 and H3K27ac, but not over the loci individ-ually marked by H3K27me3 and H3K27ac. **(D)** Feature heatmaps showing H3K4me3, H3K27ac, and CoCUT&Tag signal centered over H3K4me3-specific, shared, and H3K27ac-specific peaks. (**E**) Co-occupancy signal is concentrated at shared active sites and largely absent from H3K27ac-only distal elements. **(F)** Feature heatmaps showing H3K27me3, H3K27ac, and CoCUT&Tag signal centered over H3K27me3-specific, shared, and H3K27ac-specific peaks. **(G)** Genome annotations indicate H3K27me3-H3K27ac shared sites are enriched in blacklist-associated regions.

To test whether CoCUT&Tag preserves target specificity when two chromatin features are profiled together, we applied it to K562 cells to detect combinations of active and repressive histone marks. When either the active mark H3K4me3 or H3K27ac was paired with the repressive mark H3K27me3, the resulting CoCUT&Tag profiles remained cleanly separated, each profile mapped to its expected genomic regions with no evidence of cross-reaction from the other mark profiled in the same reaction (**Fig. 1B,C**). CoCUT&Tag profiles were also highly correlated with standard CUT&Tag for the same histone modification (**Fig. 1B**), indicating that the SunTag and MoonTag scaffolds preserve target specificity and signal quality while enabling simultaneous profiling of two chromatin features.

We next asked whether mixed-barcode fragments identify genomic regions where two chromatin features are truly linked on the same DNA molecule. As a positive control, we paired H3K4me3 and H3K27ac, two active histone marks that frequently co-occur at promoters. As expected, co-occupancy signal was enriched at shared peaks, but was much weaker at H3K4me3-specific sites and nearly absent at H3K27ac-only distal elements (DE; **Fig. 1C-E**). As a negative control, we performed CoCUT&Tag for the repressive mark H3K27me3 together with the active mark H3K27ac, which are expected to decorate largely mutually exclusive regions of the genome. In this case, shared peaks were rare and genome-wide co-occupancy signal was correspondingly sparse (**Fig. 1C,F**). The small subset of shared sites was enriched in blacklisted repetitive regions and was also marked by co-occupancy reads (**Fig. 1G**), consistent with recent evidence that H3K27me3 and H3K27ac can coexist over transposable elements in human cells^19^. Together, these results show that CoCUT&Tag distinguishes sites of true co-occupancy from genomic regions bound independently by each chromatin feature. Notably, these bulk CoCUT&Tag experiments were performed with only 25,000 cells as input, representing a substantial reduction relative to the millions of cells required for sequential ChIP and co-ChIP methods.

### Single-cell CoCUT&Tag resolves immune cell states with linked, high-precisionembeddings

We next adapted CoCUT&Tag for high-throughput single-cell profiling in peripheral blood mononuclear cells (PBMCs). We developed a combinatorial indexing workflow and compared rabbit and mouse monoclonal antibodies against H3K4me3 and H3K27me3 (**Fig. 2A;Supplementary Fig. 2A**). To identify the optimal scaffold length for single-cell profiling, we compared SunTag and MoonTag scaffolds containing 4x, 12x, or 24x antigen repeats. For both histone marks, 12x antigen repeats consistently yielded more reads per cell than 4x antigen repeats, with only a modest reduction in signal-to-noise, whereas performance declined with 24x-repeat scaffolds (**Fig. 2B-E**). We also identified rabbit and mouse monoclonal antibodies that matched or outperformed standard CUT&Tag in reads per cell when paired with 12x antigen repeats (**Supplementary Fig. 2B,C**). Thus, 12x-repeat SunTag and MoonTag scaffolds balanced signal amplification and assay quality, establishing conditions for robust single-cell CoCUT&Tag profiling.

**Figure 2.**
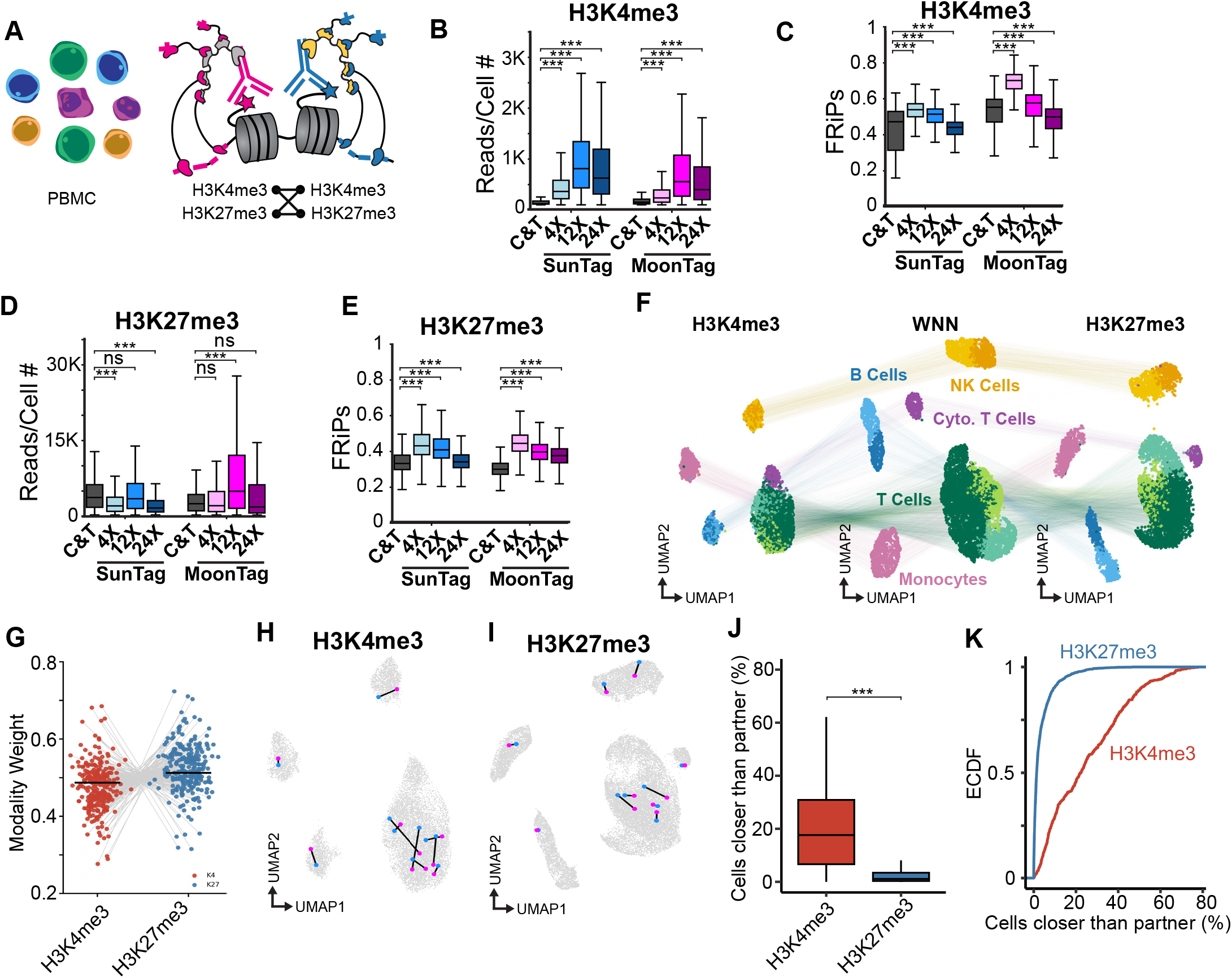
Single-cell CoCUT&Tag resolves immune cell states with linked, high-precision embeddings. **(A)** CoCUT&Tag bench-marking in PBMCs. **(B-E)** Single-cell CoCUT&Tag performance metrics for standard CUT&Tag and the indicated SunTag and Moon-Tag antigen repeat lengths. Twelve-copy antigen scaffolds improve or maintain reads per cell and library quality relative to standard CUT&Tag, whereas twenty-four-copy scaffolds reduce performance. **(F)** Weighted nearest neighbor (WNN) embedding of PBMC single-cell CoCUT&Tag data. Major immune cell populations are resolved and annotated based on cluster-specific ArchR-imputed H3K4me3 gene scores. **(G)** Relative modality weights in the WNN embedding. H3K27me3 contributes more strongly than H3K4me3 to PBMC cell-state structure in this dataset. **(H,I)** UMAP projections showing paired measurements recovered from the same cell for H3K4me3 (**H**) or H3K27me3 (**I**). Replicate measurements of the same mark map near one another in low-dimensional space. **(J,K)** Quantification of embedding fidelity between paired measurements from the same cells. The proportion of cells closer than the true partner in multidimensional space (**J**) and the cumulative distribution of this metric (**K**) show strong within-cell correspondence.

Under these conditions, paired H3K4me3 and H3K27me3 profiles from the same PBMCs resolved the expected immune populations. After quality-control filtering (**Supplementary Fig. 2D-I**), weighted nearest-neighbor analysis identified five major cell types, including B cells, NK cells, T cells, monocytes, and cytotoxic T cells, and each population was also apparent in the separate H3K4me3 and H3K27me3 embeddings (**Fig. 2F; Supplementary Fig. 2J**; Supplementary Table 1). Thus, CoCUT&Tag profiles of H3K4me3 and H3K27me3 resolve the expected immune cell states with concordant clustering between marks. In the joint embedding, H3K27me3 contributed greater modality weight than H3K4me3 (**Fig. 2G**), indicating H3K27me3 profiles captured more of the cell-type-specific chromatin variation in PBMCs.

To determine which chromatin mark produced the most precise embeddings, we profiled the same mark with two independent antibodies in the same cells (**Fig. 2H,I**). These paired profiles provide a ground-truth test of how well each chromatin mark preserves the expected proximity of the two representations of the same cell in multidimensional space. We quantified the fraction of cells lying closer to a given cell than the true replicate profile, such that lower values indicate higher precision. By this metric, H3K27me3 embeddings were more precise than H3K4me3 embeddings (**Fig. 2J,K**). Together, these results established conditions for precise single-cell CoCUT&Tag embedding of linked chromatin profiles and identified H3K27me3 as the most informative anchor for our downstream analysis of bivalent chromatin states.

### Bivalent chromatin persists in lineage-committed progenitor cells

To quantify bivalent chromatin across adult stem cells, progenitors, and differentiated cell types, we applied single-cell CoCUT&Tag to human bone marrow samples, pairing H3K27me3 with H3K4me1, H3K4me2, or H3K4me3 (**Fig. 3A**). To guide quality-control filtering of single-cell CoCUT&Tag profiles, we mixed bone marrow samples from four donors, and used souporcell for genotype-based doublet calling^20^ (**Supplementary Fig. 3**). We then removed clusters enriched for doublets and clusters with weak nucleosomal laddering, which is indicative of reduced chromatin integrity. Because H3K27me3 produced the most precise embeddings in PBMCs, we used it as the anchor for cross-modality comparison. All three CoCUT&Tag H3K4 methylation datasets recovered similar hematopoietic populations and formed coherent clusters when colored by the paired H3K27me3 assignments. Each histone mark independently resolved the hematopoietic stem cells and multipotent progenitors (HSC/MPP), and the expected lymphoid, myeloid, erythroid, dendritic, and basophil populations (**Fig. 3B;Supplementary Fig. 4A-D**). We then assigned cell type identities to clusters using ArchR-imputed H3K4me3 gene scores together with a reference single-cell RNA-seq dataset annotated by cell-surface markers^21,22^ (**Fig. 3C**).

**Figure 3.**
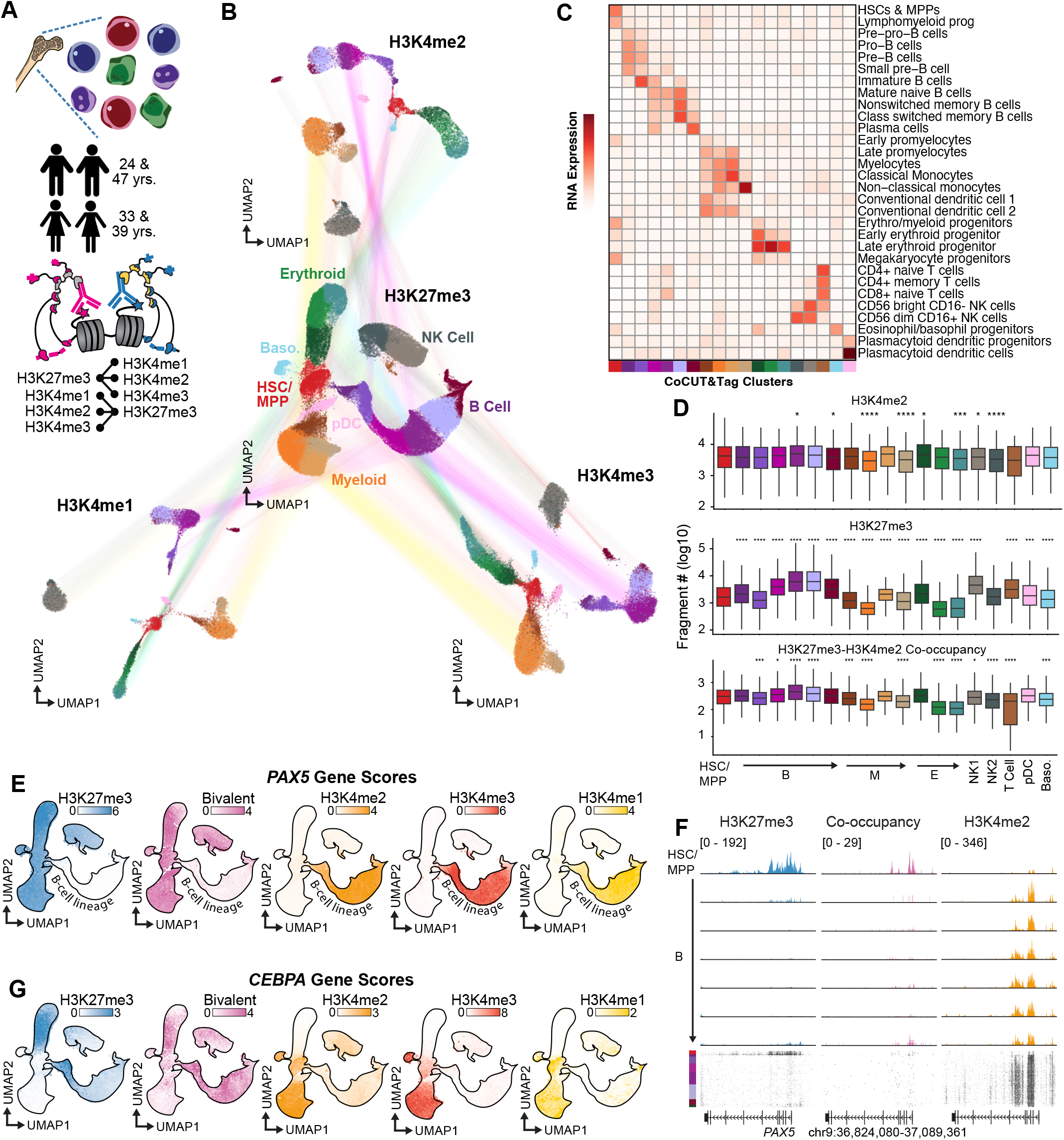
Bivalent chromatin persists in lineage-committed progenitor cells. **(A)** Bivalent chromatin was profiled in human bone marrow mononuclear cells using H3K27me3 paired with H3K4me1, H3K4me2, or H3K4me3. **(B)** Linked UMAP embeddings of H3K27me3, H3K4me2, H3K4me3, and H3K4me1 datasets, colored by H3K27me3-based cluster assignments. All four marks recover a shared hematopoietic lineage structure. **(C)** Cluster annotation based on cluster-specific ArchR-imputed H3K4me3 gene scores and expression of the corresponding genes in a reference single-cell RNA-seq dataset with surface marker-defined hematopoietic identities. **(D)** Reads per cell for H3K4me2, H3K27me3 and H3K27me3-H3K4me2 co-occupancy across hematopoietic cell states. Bivalent signal is readily detected across the hierarchy and is not maximal in HSC/MPP populations. **(E)** ArchR-imputed gene scores at *PAX5* show lineage-restricted differences in H3K4 methylation, H3K27me3, and bivalency. **(F)** Genome browser tracks across the *PAX5* locus showing H3K4me2, H3K27me3, and bivalent CoCUT&Tag signal across hematopoietic populations. Bivalent signal is detected only where both constituent marks are present. (**G**) Same as (E), but colored by the imputed gene scores for *CEBPA*.

Because CoCUT&Tag measures linked chromatin states within individual cells, we can directly quantify bivalency across cell types within the same sample. H3K4me1, H3K4me2, and H3K4me3 levels were relatively stable across hematopoietic populations, whereas H3K27me3 varied more substantially, increasing in the B-cell lineage while declining during myeloid and erythroid differentiation (**Fig. 3D;Supplementary Fig. 4E,F**). The relative abundance of each type of H3K27me3-H3K4 methylation co-occupancy reads tracked closely with H3K27me3 abundance across cell types, indicating that global variation in bivalency is driven primarily by H3K27me3. Unexpectedly, bivalent signal was not the highest in HSC/MPP cells and remained high in downstream populations, especially in the B-cell lineage (**Fig. 3D;Supplementary Fig. 4E,F**).

The persistence of bivalent signal outside HSC/MPP populations was also evident at individual genomic loci. For example, the B-lineage transcription factor *PAX5* was bivalent in HSC/MPP cells and lost H3K27me3 together with bivalent signal along the B-cell lineage as H3K4 methylation increased, consistent with resolution of bivalency during gene activation (**Fig. 3E**). Outside the B lineage, however, *PAX5* retained H3K27me3 together with detectable H3K4me2, producing strong bivalent signal in lineage-committed populations where *PAX5* remained repressed. Genome browser tracks across the *PAX5* locus showed that bivalent co-occupancy reads were concentrated at the promoter only in the cells where both marks were present, providing an internal validation that the co-occupancy signal reflects true linked chromatin states (**Fig. 3F**). By contrast, the myeloid transcription factor *CEBPA* was marked by H3K4 methylation in HSC/MPP cells without detectable bivalent signal and acquired bivalency in lineage-committed erythroid and B-cell populations (**Fig. 3G**). Thus, *PAX5* bivalency is resolved during lineage activation, whereas *CEBPA* shows that some genes become bivalent as they are developmentally repressed outside their expressed lineage. Together, our direct single-cell measurements of bivalent chromatin argue against the simple model that bivalency is uniquely concentrated in stem and multipotent progenitor cells.

### Bivalent and unmarked chromatin states distinguish classes of protein-coding genes

The abundance of total bivalent reads across cell types does not necessarily predict the number of bivalent genes because strong bivalent signal at a few loci could produce as many reads as weaker signal spread across a larger number of genes. To address this possibility, we shifted our analysis from cell-level bivalent signal to gene-level chromatin-state classification. Using Gaussian mixture modeling, we assigned thresholds for H3K27me3, H3K4me1, H3K4me2, H3K4me3, and bivalent co-occupancy across clusters (**Fig. 4A,B**). Bivalent genes showed higher H3K27me3 scores than repressed genes and higher H3K4me1 and H3K4me2 scores than active genes, but lower H3K4me3 scores (**Fig. 4C;Supplementary Fig. 5A**). These patterns indicate that bivalency is not simply an intermediate between activation and repression, but a distinct chromatin state in which activating and repressive histone marks accumulate together. Genes classified as bivalent by multiple H3K4 methylation marks showed the strongest enrichment for the Gene Ontology category “developmental process” (**Fig. 4D)**, indicating that chromatin-state assignments that are consistent across H3K4 methylation marks better distinguish biological signal from noise. To unify these chromatin-state classifications across the three H3K4 methylation profiles, we assigned each gene to a consensus active, repressed, bivalent, or unmarked state in each cluster. In HSC/MPP cells, bivalent genes were especially enriched for Gene Ontology categories related to hematopoietic lineage commitment, whereas unmarked genes were not (**Fig. 4E;Supplementary Fig. 5B**). RNA expression in HSC/MPP cells also tracked with these chromatin-state assignments, supporting the biological coherence of the classifier (**Fig. 4F**).

**Figure 4.**
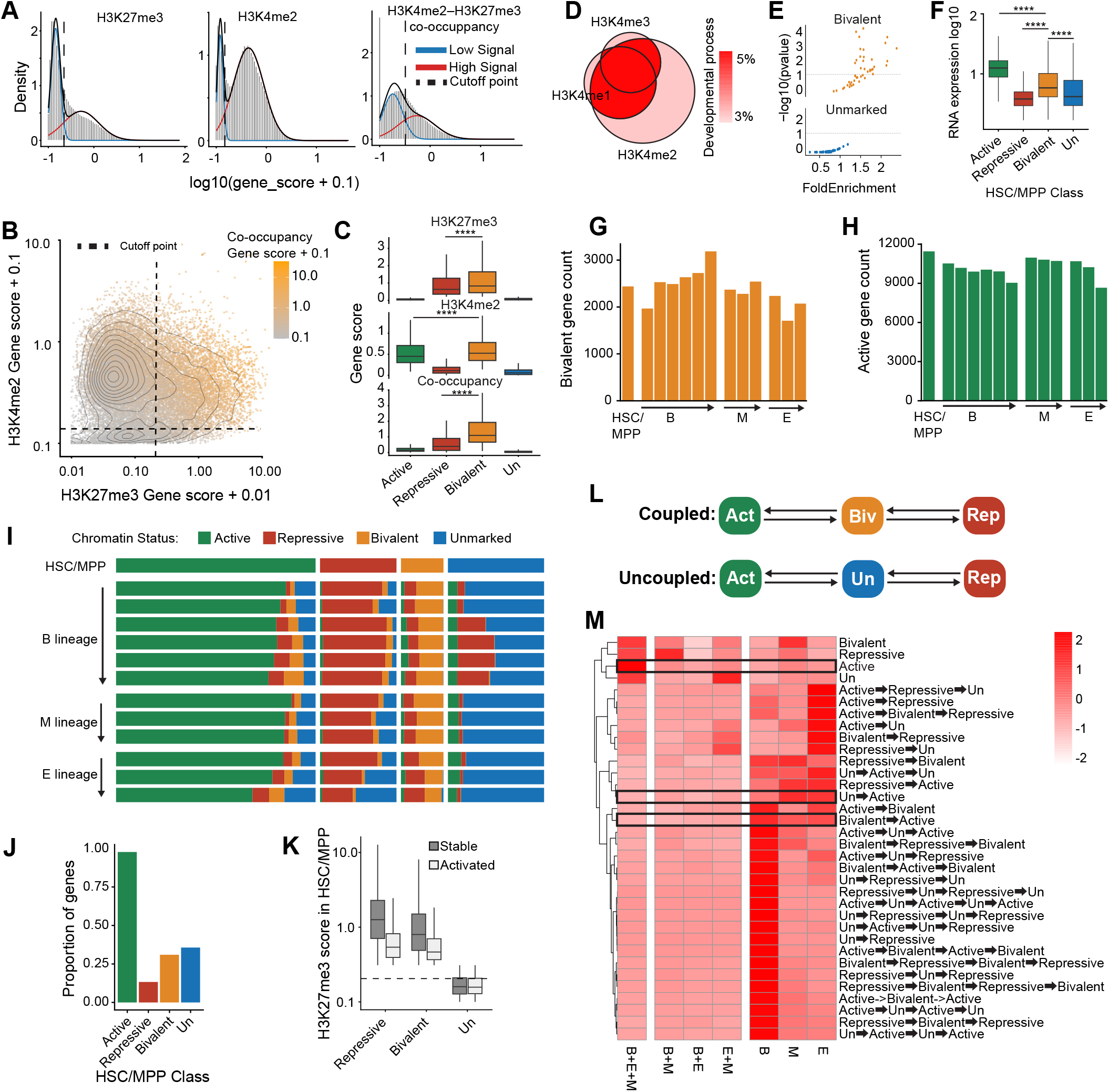
Bivalent and unmarked chromatin states distinguish classes of protein-coding genes. **(A-F)** Gene classification strategy. **(A)** Gaussian mixture modeling was used to set a threshold to distinguish genes with low and high gene scores for H3K27me3, H3K4me1, H3K4me2, H3K4me3, and bivalent co-occupancy. **(B)** Genes positive for both H3K4me2 and H3K27me3 showed the highest levels of bivalent CoCUT&Tag signal and **(C)** higher levels of H3K27me3 and H3K4me2 than the other gene classes. **(D)** Developmental gene ontology enrichment is strongest among genes with a consensus classification as bivalent across H3K4me1, H3K4me2, and H3K4me3 methylation states. **(E)** Scatter plot showing Gene Ontology enrichment for bivalent and unmarked genes in HSC/MPPs. Bivalent genes are enriched for hematopoietic developmental programs, whereas **(F)** RNA expression differs across the active, repressed, bivalent, and unmarked gene classes in HSC/MPPs. **(G)** Bar graph of the absolute number of consensus bivalent genes or **(H)** consensus active genes in the HSC/MPP and lineage-committed B, M, and E clusters. **(I)** Fate map of genes beginning in active, repressed, bivalent, or unmarked states in HSC/MPPs and transitioning through the B, M, and E lineages. **(J)** Proportion of genes from each initial HSC/MPP chromatin state that transition to an active state in at least one downstream cluster. **(K)** H3K27me3 levels in HSC/MPPs for genes that do or do not transition to active states. Genes activated from bivalent and repressed states begin with substantial H3K27me3, whereas genes activated from unmarked states do not. **(L)** Model for the two regulatory identities of coding genes. Polycomb/Trithorax-coupled genes transition between bivalent and repressive or active chromatin states, and uncoupled genes transition between unmarked and repressive or active states. **(M)** Heatmap showing the lineage specificity of gene-state transitions. Active genes tend to remain active across multiple lineages, whereas bivalent-to-active and unmarked-to-active transitions are predominantly lineage restricted. Unmarked-to-repressed transitions are especially enriched in the B lineage.

Gene-level classification confirmed that HSC/MPP cells do not contain the largest set of bivalent genes. Instead, the number of bivalent genes increased in the B-cell and myeloid lineages and declined in the erythroid lineage (**Fig. 4G**). In contrast to the bivalent genes, the number of active genes was highest in HSC/MPP cells and decreased across all three lineages (**Fig. 4G,H**). Thus, HSC/MPPs are characterized by a more active chromatin landscape that was apparent at the gene level, but not from the global abundance of H3K4 methylation marks.

We next asked how the chromatin status of bivalent, repressed, active, and unmarked genes classified in HSC/MPP cells changed in downstream lineages. Most bivalent genes remained bivalent or shifted to a repressed state, and only a minority of bivalent genes became active in any given lineage, and this fraction was slightly lower than that for unmarked genes (**Fig. 4I,J**). Even fewer repressed genes became active, and this could not be explained simply due to marginal H3K27me3 calls, because repressed genes that underwent repressed-to-active transitions had H3K27me3 levels in HSC/MPP cells comparable to those of activated bivalent genes (**Fig. 4K**). Strikingly, almost no genes transitioned between the bivalent and unmarked states in either direction (**Fig. 4I**), but many unmarked genes in the B-cell lineage did transition to a repressed state (**Supplementary Fig. 5C**). These results suggest that bivalent and unmarked states are not merely temporary regulatory configurations but instead represent two fundamental chromatin states that rarely interconvert with one another and can each resolve toward active or repressed states (**Fig. 4L**).

We then asked whether chromatin-state changes were shared across lineages or instead reflected activation of different genes in each developmental branch. Active genes in HSC/MPP cells usually remained active across multiple downstream lineages (**Fig. 4M**). In contrast, most bivalent or unmarked genes became active in only one lineage. In the rare event that repressed genes transitioned to an active state it was also typically restricted to a single lineage (**Fig. 4M**). Thus, activation of developmentally regulated genes is generally lineage specific, with distinct gene sets activated in different lineages rather than shared across many lineages. Together, these data show that bivalent chromatin is a specialized regulatory state that persists during lineage commitment and differentiation, and is more readily activated than fully repressed chromatin.

### Bivalent genes reach higher levels of active chromatin marks and gene expression than unmarked genes

Although bivalent and unmarked genes had a similar overall propensity to become active during lineage commitment, they may nevertheless do so with different activation dynamics. To test this, we ordered cells along B-, myeloid-, and erythroid-lineage pseudotime trajectories (Fig. 5A). We split the HSC/MPP cluster into two subgroups and found that one already showed active marks at genes associated with the myeloid lineage, indicating a potential myeloid bias. We used the more stem-like subgroup as the trajectory root (Supplementary Fig. 6A-D). Bivalent genes that transitioned to an active state showed the expected loss of H3K27me3 and bivalent co-occupancy together with gains in H3K4 methylation along pseudotime (**Fig. 5B;Supplementary Fig. 6E-G**). However, when trajectories were scaled to compare their relative progression, bivalent and unmarked genes that became active showed nearly identical timing of H3K4 accumulation across all three lineages (**Fig. 5C-F;Supplementary Fig. 7A-C**). By contrast, the absolute magnitude of H3K4 accumulation was substantially higher at bivalent genes than at unmarked genes, and this pattern was evident across multiple H3K4 methylation marks (**Fig. 5B;Supplementary Fig. 6E-G**). Thus, bivalent genes do not gain active chromatin earlier than unmarked genes but instead reach stronger active chromatin signal upon induction.

**Figure 5.**
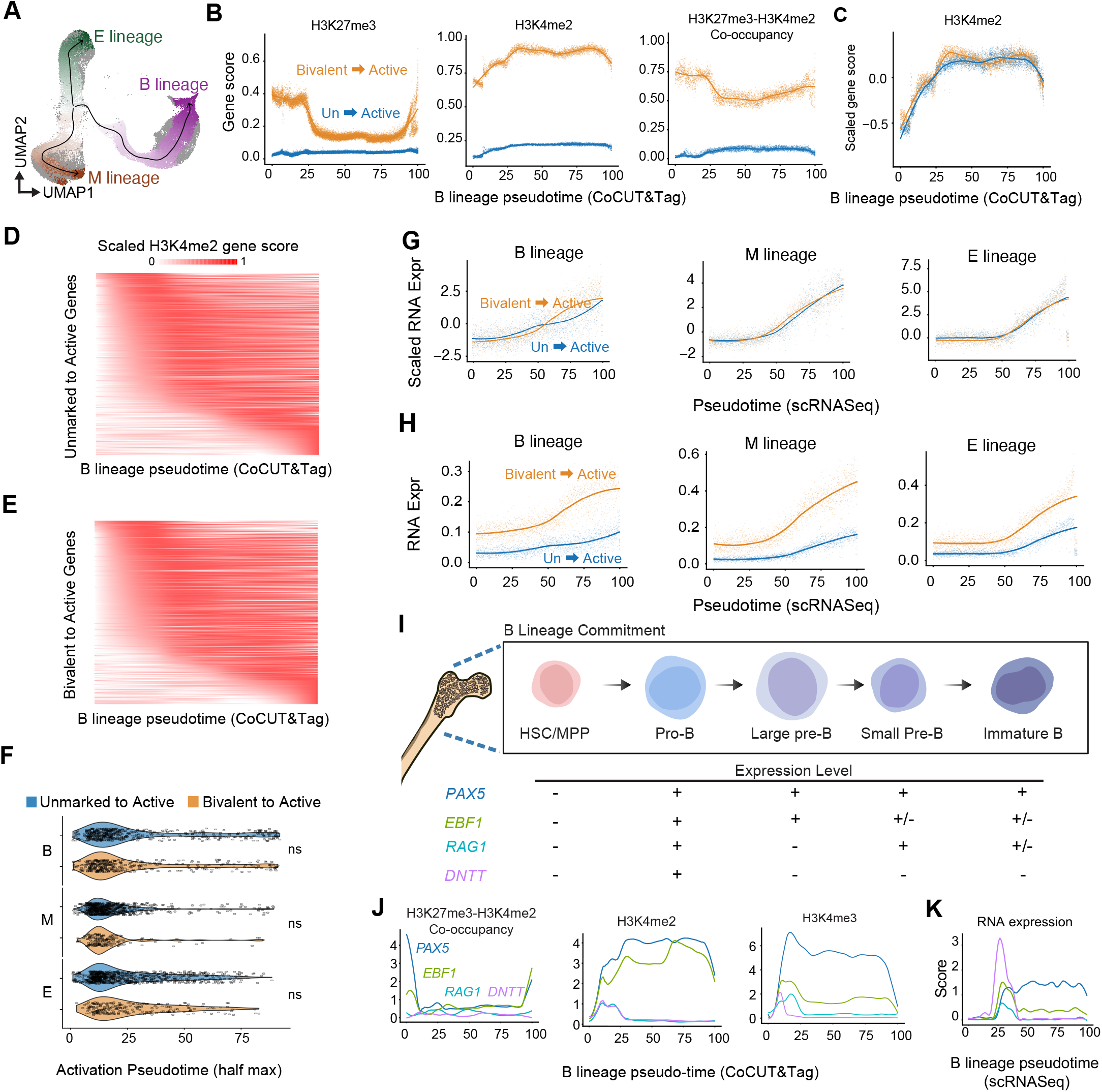
Bivalent genes reach higher levels of active chromatin marks and gene expression than unmarked genes. **(A)** H3K27me3 UMAP showing the inferred B, E and M-lineage pseudotime trajectories. **(B)** Smoothed CoCUT&Tag gene scores for H3K27me3, H3K4me2, and H3K27me3-H3K4me2 co-occupancy across B-lineage pseudotime for genes that transition from bivalent-to-active chromatin (orange) or from unmarked-to-active chromatin (blue). **(C)** Scaling the H3K4me2 gene scores from (B) shows that unmarked and bivalent genes gain H3K4me2 with similar timing. **(D)** Heatmap comparing the H3K4me2 gene scores across B lineage pseudotime for the unmarked genes (top) versus the bivalent genes (bottom) that transitioned to an active state. Genes are rank ordered according to the point they reach the half max H3K4me2 gene scores in pseudotime. **(F)** Violin plot showing the H3K4me2 activation timing (half-max point in pseudotime) is similar for bivalent and unmarked genes across the B, M and E lineages. **(G,H)** Average RNA expression across pseudotime for genes that transition from bivalent-to-active chromatin or from unmarked-to-active chromatin in the B, M, and E lineages, shown as scaled expression (G) or unscaled expression (H). Bivalent genes activate with similar timing but reach higher expression levels. **(I)** Schematic of B-cell differentiation stages represented along the pseudotime trajectory. **(J)** CoCUT&Tag gene scores for representative B-lineage regulators across B-lineage pseudotime. *PAX5* and ***E****BF1* lose H3K27me3-H3K4me2 co-occupancy signal as H3K4me2 and H3K4me3 increase, whereas *RAG1*, and *DNTT* gain H3K4me2 with little or no bivalent signal. **(K)** Corresponding scRNA-seq expression trajectories for the genes shown in (I). *DNTT* is not bivalent but activates with high amplitude during B-cell differentiation.

We next asked whether this increased signal amplitude was also reflected in RNA expression. Among genes activated during differentiation, bivalent genes were more likely than unmarked genes to show greater than twofold induction relative to HSC/MPP cells across the B-, myeloid-, and erythroid-lineage trajectories (**Supplementary Fig. 7D**). Bivalent genes and unmarked genes again showed similar timing of activation, but once activated the bivalent genes reached substantially higher RNA expression levels (**Fig. 5G,H**). Together, these results indicate that bivalent chromatin is not associated with earlier gene induction, but instead with stronger chromatin activation and higher gene expression.

To illustrate how chromatin and expression dynamics are coupled at individual loci, we examined representative genes expressed at distinct stages of B-cell differentiation. The B-lineage transcription factors *PAX5* and *EBF1* lost bivalent signal as H3K4me2 increased, consistent with activation from bivalent promoters (**Fig. 5I,J**). By contrast, *RAG1* and *DNTT*, which direct programmed mutagenesis of the B-cell receptor, showed distinct regulatory patterns. The *DNTT* locus was not bivalent, yet underwent a rapid burst of expression, illustrating that high-amplitude gene expression can also arise from non-bivalent chromatin states (**Fig. 5K**). Notably, although both *DNTT* and *RAG1* showed two peaks of H3K4me2 signal, *DNTT* exhibited only a single peak of H3K4me3 that coincided with the first peak of H3K4me3 on the *RAG1* gene (**Fig. 5J**). This mirrors previous studies showing that *DNTT* expression spikes early during B-cell commitment, whereas *RAG1* is activated early and then reactivated later^23,24^. These examples show that CoCUT&Tag can resolve fine temporal patterns of gene regulation.

### Bivalent gene activation is associated with more numerous and stronger enhancer elements

Bivalent and unmarked genes may also differ in the organization of the cis-regulatory elements that drive their distinct activation patterns. To test this, we defined candidate cis-regulatory elements from H3K4me2 peaks across the B-, myeloid-, and erythroid-lineage trajectories and quantified H3K4me2, H3K27me3, bivalency, H3K4me3, and H3K4me1 across the corresponding genomic windows (**Fig. 6A**). Dimensionality reduction separated these elements primarily into two large groups, one marked by H3K4me3 and the other by H3K4me1 (**Fig. 6B,C**). H3K4me3-marked elements were enriched near gene transcription start sites (TSSs) and were therefore classified as promoter elements, whereas H3K4me1 was enriched at distal elements (**Supplementary Fig. 8A,B**). Because H3K4me1 is associated with enhancers that activate nearby promoters^25^, we consider these distal H3K4me1-marked peaks as enhancer elements.

**Figure 6.**
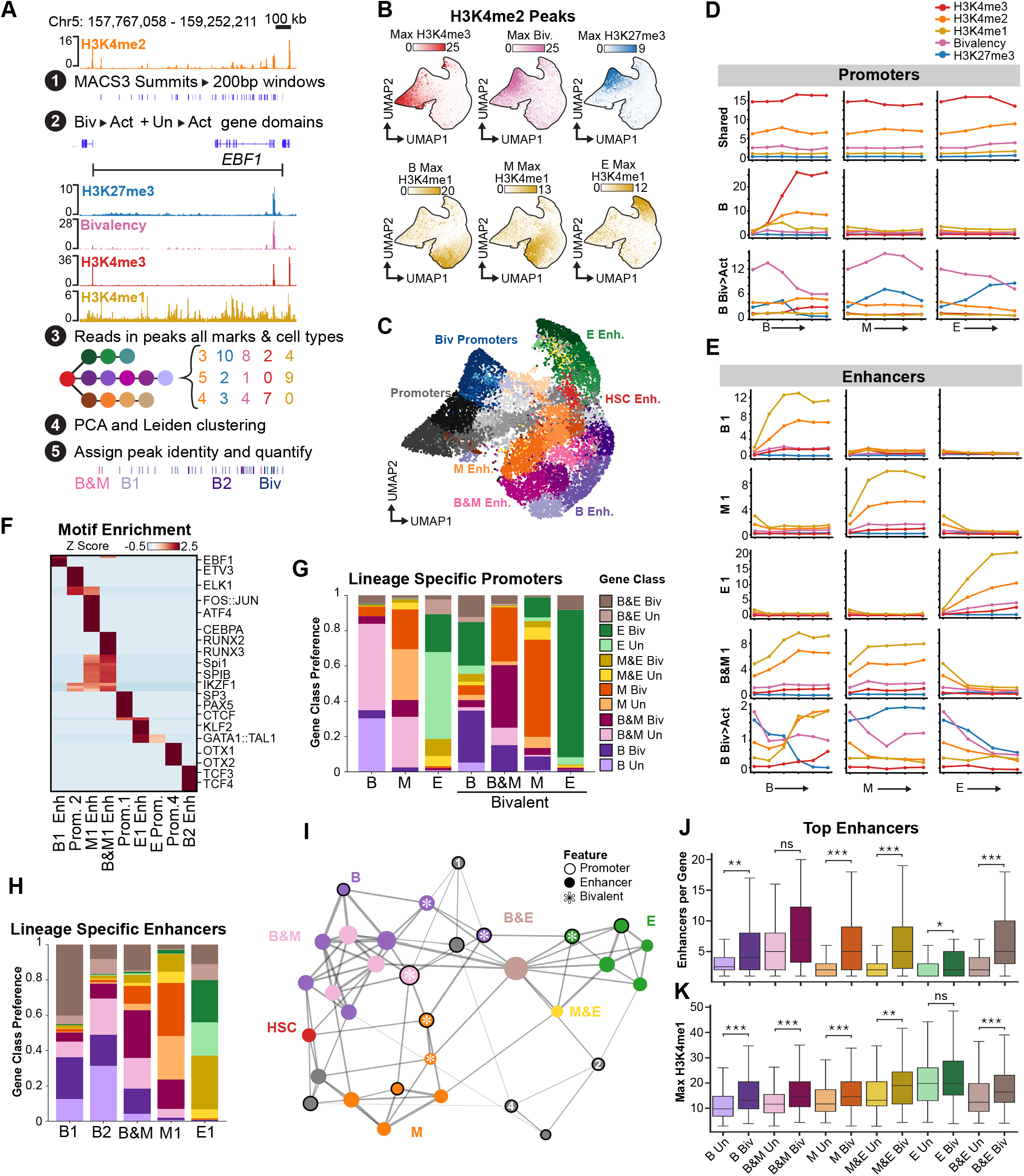
Bivalent gene activation is associated with more numerous and stronger enhancer elements. **(A)** Workflow for regulatory element analysis. H3K4me2 peaks were used to define 200-bp regulatory elements associated with genes that transitioned from bivalent to active or from unmarked to active states. H3K4me2, H3K27me3, bivalency, H3K4me3, and H3K4me1 signal were quantified across hematopoietic clusters and used for dimensionality reduction and Leiden clustering. **(B)** UMAP of H3K4me2-defined regulatory elements colored by weighted chromatin feature values. **(C)** Regulatory element classes inferred from chromatin features and genomic position. H3K4me3-high clusters correspond to promoters, whereas lineage-specific H3K4me1-high clusters correspond to distal enhancers. Bivalent elements are predominantly promoter associated. **(D)** Average chromatin profiles for promoter classes across the B, M, and E lineages. Promoter classes include constitutive promoters, lineage-specific promoters that gain H3K4me3 *de novo*, and bivalent promoters that resolve through loss of H3K27me3 together with gains in H3K4me3. **(E)** Average chromatin profiles for enhancer classes across the B, M, and E lineages. Enhancer classes include lineage-specific B, M, and E enhancers, enhancer groups shared across two lineages, and rare bivalent enhancers. **(F)** Motif enrichment across promoter and enhancer classes. Lineage-specific enhancers are enriched for expected lineage regulators, including EBF1 and TCF3 in B-lineage enhancers, CEBP and AP-1 motifs in myeloid enhancers, RUNX, SPI and IKZF motifs in shared B/myeloid enhancers, and GATA1::TAL1 in erythroid enhancers. **(G)** Enrichment of promoter classes across genes undergoing lineage-specific unmarked-to-active or bivalent-to-active transitions. Lineage-specific promoters that gain H3K4me3 *de novo* are preferentially associated with unmarked-to-active genes, whereas bivalent promoters are enriched at bivalent-to-active genes. **(H)** Enrichment of enhancer classes across the same gene categories. Lineage-specific enhancers occur in both unmarked-to-active and bivalent-to-active genes, with modest lineage-dependent biases. **(I)** Network showing co-occurrence of regulatory element classes within gene domains. Lineage-specific promoters and enhancers group together with shared lineage elements organized in between the corresponding lineage-specific elements. **(J,K)** Quantification of enhancer number per gene (J) and enhancer H3K4me1 signal (K) for top lineage-associated enhancer classes. Genes activated from bivalent chromatin carry more enhancers with stronger H3K4me1 signal than genes activated from unmarked chromatin.

Promoter elements were organized along opposing H3K4me3 and H3K27me3 gradients, with the highest bivalency at elements occupying the middle positions of the promoter clusters in UMAP space (**Fig. 6C,D;Supplementary Fig. 8C–F**). By contrast, enhancers were organized by lineage-specific H3K4me1 accumulation and formed groups enriched in the B, myeloid, or erythroid lineages, together with smaller groups shared between B and myeloid (B&M), B and erythroid (B&E) and erythroid and myeloid (E&M) lineages (**Fig. 6C,E;Supplementary Fig. 8G**). We did not identify promoters or enhancer clusters that gained H3K4 methylation across all three lineages, providing further support for the model that differentiation is driven by multiple distinct lineage-specific programs rather than a single universal program. Among these regulatory elements, 21% (1,177/5,636) of promoter elements were classified as bivalent, and were marked by lineage-specific gains in H3K4me3 together with reductions in H3K27me3. In comparison, 6.6% (584/8,875) of the enhancer elements we identified showed gains in H3K4me1 and reductions in H3K27me3, and were classified as bivalent. These results show that bivalent chromatin is enriched over promoters.

Motif analysis supported the lineage-specific assignments of these regulatory elements and further distinguished enhancers from promoters (**Fig. 6F**; Supplementary Table 2). Enhancers with the strongest lineage-specific H3K4me1 signal were enriched for the expected transcription factor motifs, including EBF1 and TCF3 in B-lineage enhancers, CEBP and AP-1 family motifs in myeloid enhancers, SPI1 and RUNX motifs in shared B&M enhancers, and GATA1::TAL1 in erythroid enhancers (**Fig. 6F;Supplementary Fig. 8H**; Supplementary Table 2). In comparison, lineage-specific promoters showed weaker motif enrichment than enhancers, but the shared promoters (Prom. 1, Prom. 2 and Prom. 4) that retained high H3K4me3 levels across lineages were enriched for motifs such as ELK1, ETV3, SP/KLF, PAX5, and CTCF (**Fig. 6F**; Supplementary Table 2). Together, these results indicate that the regulatory information specifying lineage-specific gene activation resides primarily in enhancer elements rather than promoters.

We next compared how different groups of regulatory elements were distributed for bivalent versus unmarked genes undergoing activation. As expected, promoters that gained H3K4me3 *de novo* were enriched near unmarked genes, whereas bivalent promoters were enriched near bivalent genes (**Fig. 6G**). In contrast, enhancer elements were distributed across both gene classes (**Fig. 6H**). A co-occurrence network showed that promoter and enhancer groups were organized into lineage-biased combinations, with regulatory elements shared across two lineages occupying intermediate positions between lineage-specific groups (**Fig. 6I**).

We next compared the number and H3K4me1 signal intensity of regulatory elements found in bivalent and unmarked gene domains, defined as the intervals between the neighboring upstream and downstream genes. Among the top lineage-specific enhancers with the strongest H3K4me1 signal, bivalent genes were linked to more enhancer elements per gene than unmarked genes, and these bivalent gene-linked enhancers also reached higher H3K4me1 signal (**Fig. 6J,K**). The same trend was observed when all enhancer classes were included, indicating that the effect was not driven by a particular subset definition (**Supplementary Fig. 9C,D**). Bivalent genes were also associated with more promoter elements than unmarked genes, but those promoters did not show higher H3K4me3 signal (**Supplementary Fig. 9E,F**). Thus, bivalent and unmarked genes differed both in promoter chromatin state and in the number and strength of associated enhancers, with bivalent genes linked to more numerous and stronger enhancers.

### Polycomb restraint raises the enhancer threshold for bivalent gene activation

We next tested the hypothesis that Polycomb-associated H3K27me3 at promoters raises the enhancer input required for activation of bivalent genes. This model makes two predictions. First, bivalent genes should remain transcriptionally silent despite the accumulation of H3K4me1 at nearby enhancer elements, whereas unmarked genes should become active with fewer such enhancer elements or lower H3K4me1 signal. Second, reducing Polycomb restraint should preferentially activate bivalent genes. To test these predictions, we focused on the erythroid lineage, where lineage-specific enhancer classes were most clearly resolved.

Using the same H3K4me2-based peak-calling strategy but applied to all lineages and gene domains, we identified the full set of enhancers that gained H3K4me1 specifically in the erythroid lineage (**Supplementary Fig. 10A,B**). As expected from the previous enhancer-per-gene-domain analysis, bivalent gene activation was associated with more erythroid enhancer elements per gene and with higher H3K4me1 signal than unmarked genes (**Fig. 7A,B**). Strikingly, however, some genes that remained bivalent (E. Biv) still had substantial erythroid enhancer signal, often exceeding the number of enhancers associated with unmarked genes (**Fig. 7A,B**). *GATA5* provides one such example, and remained bivalent despite nearby erythroid enhancers that showed strong H3K4me1 signal (**Fig. 7C**). The same conclusion held when all putative erythroid enhancer classes were included rather than only the strongest groups (**Supplementary Fig. 10C,D**). The hemoglobin locus illustrates the activated end of this spectrum, with a large set of nearby erythroid enhancers and a transition from bivalent to active during erythroid differentiation (**Fig. 7D**). Thus, bivalent genes can remain transcriptionally silent despite nearby lineage-associated enhancers, consistent with a higher activation threshold imposed by Polycomb-associated chromatin.

**Figure 7.**
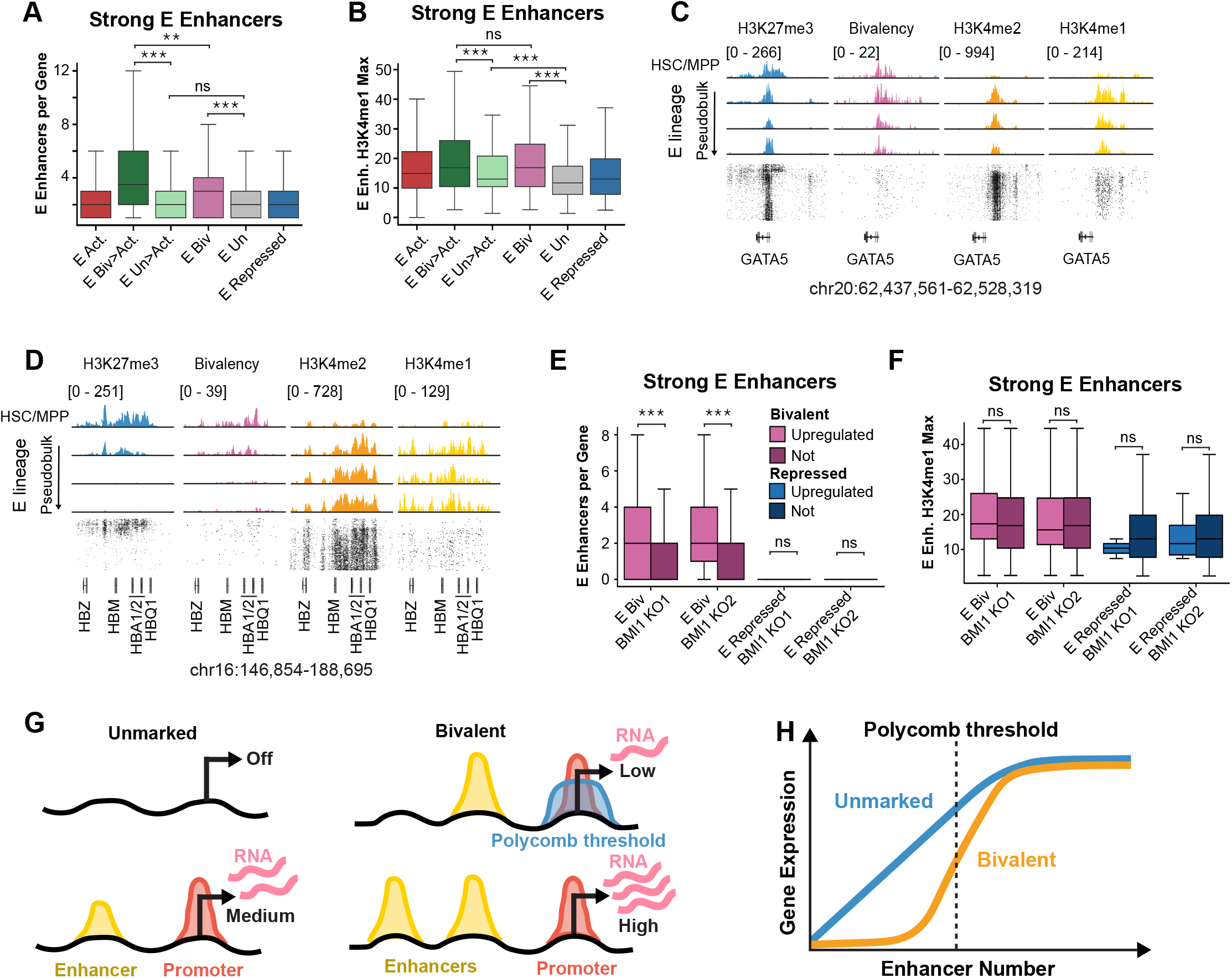
Polycomb restraint raises the enhancer threshold for bivalent gene activation. **(A,B)** Number of strong erythroid enhancers per gene (A) and maximal enhancer H3K4me1 signal (B) for genes that transition from bivalent to active, remain bivalent, transition from unmarked to active, remain unmarked, or remain repressed during erythroid differentiation. Bivalent genes that remain inactive already carry substantial erythroid enhancer input, whereas genes that transition from bivalent to active are associated with stronger enhancers. **(C)** Genome browser tracks across the *GATA5* locus showing H3K27me3, H3K27me3-H3K4me2 co-occupancy, H3K4me2, and H3K4me1 across erythroid pseudotime. *GATA5* remains bivalent despite association with multiple strong erythroid enhancers. **(D)** Same as (C) but showing the hemoglobin locus which carries a large repertoire of erythroid enhancers and transitions from bivalent to active during erythroid differentiation. **(E,F)** Analysis of *BMI1* knockout RNA-seq data from primary HSCs differentiated toward the erythroid lineage. Among genes classified as bivalent throughout the erythroid lineage, those upregulated after *BMI1* loss are enriched for greater numbers of strong erythroid enhancers per gene (**E**), whereas enhancer H3K4me1 signal is more similar between upregulated and non-upregulated bivalent genes (**F**). The same enrichment is not observed for genes in the repressed state. **(G)** Schematic showing enhancer-dependent activation of unmarked and bivalent genes. Polycomb restraint of bivalent promoters increases the demand for nearby lineage-associated enhancers, delaying high-level expression until sufficient activating input accumulates. Blue = H3K27me3; Red = H3K4me3; Yellow = H3K4me1. **(H)** Threshold model relating enhancer number to gene expression. Unmarked genes show a more direct response to enhancer activation, whereas bivalent genes are buffered against enhancer activation, producing a delayed, switch-like rise in expression.

We next asked whether bivalent genes are preferentially activated when Polycomb function is reduced. Using public datasets from CRISPR-mediated deletion from human erythroid progenitors^26^, we found that the subset of bivalent genes that were upregulated after BMI1 knockout included *GATA5* and were associated with more erythroid enhancers per gene than bivalent genes that were not upregulated (**Fig. 7E;Supplementary Fig. 10E**). This enrichment was reproducible across gRNAs, but was not observed among the repressed genes that were upregulated, indicating that bivalent genes were specifically sensitive to Polycomb-mediated repression. By contrast, H3K4me1 signal did not differ significantly between upregulated and non-upregulated bivalent genes (**Fig. 7F;Supplementary Fig. 10F**). In the intact system, however, both enhancer number and enhancer-associated H3K4me1 signal distinguished bivalent genes that became active from those that remained bivalent (**Fig. 7A,B**). Thus, when Polycomb restraint is reduced, the number of nearby enhancers predicts derepression better than enhancer-associated H3K4me1 signal alone.

Together, these results support a threshold model in which Polycomb restraint keeps bivalent genes silent despite the buildup of activating input from nearby lineage-associated enhancers (**Fig. 7G**). In this model, expression of unmarked genes increases more gradually with enhancer input, whereas bivalent genes are buffered against enhancer-driven activation until a Polycomb-dependent threshold is crossed, after which high-level expression is rapidly achieved (**Fig. 7G,H**).

## Discussion

The CoCUT&Tag assay developed here overcomes the key challenges associated with multifactorial CUT&Tag-based assays. CUT&Tag depends on recruiting multiple Tn5 complexes to each chromatin site and on the insertion of two correctly oriented DNA adapters for PCR amplification, which is achieved through secondary-antibody amplification^27,28^. However, using conventional protein A-based tethering methods for joint profiling is subject to cross-reaction, because protein A-based tethering cannot distinguish between antibodies. To solve this specificity problem while preserving local amplification, we replaced the secondary antibodies with anti-rabbit and anti-mouse nanobodies fused to the SunTag and MoonTag antigen-peptide repeats, and replaced protein A-Tn5 with anti-SunTag-Tn5 and anti-MoonTag-Tn5. This nanobody-scaffold-based design keeps the simple workflow and sensitivity of standard CUT&Tag intact, while enabling recovery of single-target reads together with mixed-barcode co-occupancy reads from low-input bulk samples and high-throughput single-cell formats.

We revisited the model that bivalent chromatin is associated with a multipotent stem-cell state. To our knowledge, we generated the first single-cell maps of directly linked H3K27me3 and H3K4 methylation states in an adult stem cell system. Within the HSC/MPP population, we did not observe elevated levels of bivalency by either reads-per-cell or gene-level analysis, which argues against a model where the levels of bivalent chromatin correspond to the degree of stem-cell multipotency. Instead, we found that bivalency persists during hematopoiesis and increases in several lineage-restricted cell types, reaching especially high levels in the B-cell lineage. This conclusion differs from earlier promoter-centered miniChIP-on-chip studies which indicated that HSCs carry an especially high number of bivalent promoters^29^. Several differences in study design may help explain this discrepancy. First, the earlier comparisons were specific to profiles of pre-megakaryocyte erythroid progenitors and T cells, whereas we observed the most substantial accumulation of bivalency in the myeloid and B-cell lineages. Second, the prior studies used highly purified murine HSCs, whereas we did not distinguish between human HSCs and MPPs. However, the global changes in bivalency and other chromatin states that we observed likely extend to human HSCs because we did not observe a rare subset of cells marked by exceptionally high bivalency in the HSC/MPP compartment. Most importantly, previous studies inferred bivalency from the overlap of H3K4me3 and H3K27me3 profiles generated in separate samples, whereas our classification required H3K4 methylation, H3K27me3, and directly measured co-occupancy reads to exceed threshold within the same single-cell profiles.

In contrast, we did observe that HSC/MPP cells retain the largest proportion of genes in an active chromatin state. This is consistent with models suggesting that stem and progenitor cells maintain a relatively permissive chromatin landscape and elevated transcriptional capacity^30^. This observation may also be related to recent work showing that adult multipotent stem and progenitor cells can occupy a “hypertranscription” state that is defined by elevated levels of global gene expression^31,32^. Consistent with this broader active chromatin landscape, our analysis of chromatin-state transitions and enhancer elements suggests that differentiation is accompanied by a narrowing of active gene programs toward lineage-specific regulatory states. More broadly, pluripotent stem cells are characterized by a more open and dynamic chromatin landscape^33^, and also undergo periods of hypertranscription^34^. Thus, rather than being uniquely defined by bivalency, HSCs and MPPs may be more accurately characterized by a globally elevated number of genes maintained in an active chromatin configuration, with bivalency marking a more restricted subset of developmentally regulated genes.

Our results also help refine the model that bivalent chromatin maintains developmental genes in a poised state. CoCUT&Tag data are compatible with single-cell pseudotime analysis, which allowed us to directly test how bivalency is related to the kinetics of gene activation. We found that bivalent and unmarked genes show similar timing of H3K4 accumulation and increases in RNA expression across the myeloid, erythroid, and B-cell lineages. These results argue against the idea that bivalency confers a simple kinetic advantage for activation. This is consistent with prior studies of embryonic stem cell differentiation into extra-embryonic tissues, where bivalent genes are also activated at the same time as unmarked genes^11^. Rather, we find that the key distinction between bivalent and unmarked genes lies in the magnitude of activation. Bivalent genes acquired stronger active chromatin signals, were more likely to show greater than twofold increases in gene expression, and on average reached higher RNA expression levels than unmarked genes. Outside of this kinetic model, we did find evidence supporting the role of bivalent chromatin as a poised state. Bivalent genes were more likely than fully repressed genes to transition to an active state during lineage commitment, indicating that they are poised not for faster activation, but because they require less additional enhancer input for activation than genes in a purely repressed state.

By comparing the accumulation of H3K4 methylation and H3K27me3 over regulatory elements we identified differences in cis regulatory configurations that underlie the propensity for high-magnitude activation of bivalent genes. Bivalent genes were linked to more numerous and stronger lineage-associated enhancers than unmarked genes, yet some remained silent despite substantial nearby enhancer activity. We therefore propose an enhancer-threshold model in which Polycomb-associated H3K27me3 raises the regulatory input required for productive activation. In this view, bivalency shifts the activation curve, allowing lineage-associated enhancers to accumulate and, in some cases, exert low-level transcriptional activation without immediately triggering high gene expression. Then, once sufficient activation of nearby enhancers is reached, transcription proceeds with high amplitude. The preferential derepression of enhancer-rich bivalent genes following *BMI1* loss strongly supports this interpretation and is consistent with prior evidence that tissue-specific bivalent loci are particularly sensitive to disruption of Polycomb repression^15^. Thus, Polycomb-mediated repression may act as a buffer against premature or weak enhancer-driven activation. This model also fits with growing evidence that enhancers act combinatorially^35^, such that individual enhancers contribute subthreshold regulatory input that becomes productive only when additional synergistic enhancers are gained during lineage progression. This framework also helps to reconcile classical models of bivalency as a poised state with perturbation studies showing that Polycomb and Trithorax activities oppose one another during hematopoietic differentiation^9,10,12^.

In our revised model, bivalency constitutes a distinct regulatory state rather than acting as a simple intermediate between activation and repression. This is supported by our observations that bivalent genes carried higher H3K27me3 and H3K4me1 and H3K4me2 than either repressed or unmarked genes, and that bivalent and unmarked genes showed very little interconversion. Importantly, we did observe that both bivalent and unmarked genes transition between active or repressed states. This implies that the bivalent chromatin state is genetically encoded at a defined subset of developmental loci rather than arising transiently at any silent promoter. In vertebrates, Polycomb recruitment has been linked to CpG-rich promoters^36,37^, and the Polycomb group protein MTF2 binds unmethylated CpG islands^38^. In this context, it is notable that we identified a promoter class enriched among bivalent genes that also showed strong enrichment for SP/KLF-family motifs, including SP3, the closest vertebrate homolog of the Sp1-like factor for pairing sensitive silencing (Spps). In *Drosophila*, Polycomb is targeted to developmental genes by Polycomb response elements that are recognized by DNA-binding factors including Pleiohomeotic and Spps^39-42^. Analogous Polycomb response elements have been difficult to define in vertebrates. However, our data support a model that protein-coding genes in humans can be broadly separated into two regulatory identities: genes where the function of Polycomb and Trithorax group proteins are coupled and those where they are uncoupled. Defining the sequence features that shape Polycomb and Trithorax responsiveness and how they establish stable coupled versus uncoupled gene classes across development will be an important direction for future work.

This enhancer-threshold model may also explain why Polycomb and Trithorax activity is often linked at genes encoding potent developmental regulators. By raising the threshold for productive activation, Polycomb restraint can shield these developmental genes from weak or unwanted activation by nearby enhancers. From an evolutionary perspective, this may help assemble lineage-specific regulatory landscapes by allowing cells to accumulate partial regulatory input and sample intermediate states without triggering full transcriptional activation. Weak enhancer signals would therefore remain latent unless reinforced by additional synergistic elements, at which point activation could proceed with high amplitude. Thus bivalent chromatin may preserve developmental flexibility while preventing rewiring of developmental gene expression by enhancer co-option.

Our study highlights several important questions related to the function of bivalent chromatin. We suspect this enhancer-threshold framework applies broadly across tissues and organisms, but it will be important to test that idea directly by applying single-cell CoCUT&Tag to other developmental systems. For example, by extending CoCUT&Tag to endodermal transitions, hemogenic endothelium, and the emergence of blood stem cells it will be possible to define how tissue-specific patterns of bivalency are established. The association between enhancer architecture, bivalent chromatin, and BMI1-sensitive derepression strongly supports a bivalency threshold model, but direct perturbation of enhancer number, enhancer strength, and Polycomb activity at individual loci is needed to establish the rules governing activation from bivalent and unmarked chromatin states. In addition, applying CoCUT&Tag to profile additional chromatin regulators, developmental systems, and disease contexts should make it possible to define how linked chromatin states encode other types of regulatory potential.

## Resource availability

### Lead Contact

Questions related to this study and requests for resources and reagents should be directed to the lead contact, Derek Janssens, and will be fulfilled in a timely manner.

### Materials Availability

Materials generated in this study will be available upon reasonable request. The completed pTBX1-αRabbit-12xSunTag-sfGFP (Addgene plasmid # 254519), pTBX1-αMouse-12xMoonTag-mCherry2 (Addgene plasmid # 254520), pTBX1-αSunTag-Tn5 (Addgene plasmid # 254521) and pTBX1-αMoonTag-Tn5 (Addgene plasmid # 254522) constructs are available through Addgene.

### Data and Code Availability

The bulk and single-cell CoCUT&Tag primary sequencing data as well as the bulk and single-cell CUT&Tag primary sequencing data are available through the Gene Expression Omnibus under the accession numbers: GSE327819, GSE327821, GSE327816 and GSE327824. The hg38 aligned and processed single-cell data complete with cell specific barcodes and metadata are available through the zenodo data sharing portal^43^. All custom code used for data analysis and figure generation are also available through zenodo^44^. Any additional information required to reanalyze the data reported in this study will be made available upon reasonable request made to the lead contact.

## Acknowledgements

We thank Dr. Sarah Leichter for suggestions regarding the construction of the SunTag and MoonTag scaffold design, as well as Terri Bryson, Yiling Xu and Dr. Steve Henikoff for help assembling necessary reagents for the project. We thank Drs. Weifang Wu, Yvonne Fonduffe-Mittendorf, Connie Krawczyk, Peter Laird, Johnathan Licht and Darrell Chandler as well as all members of the Janssens lab and the rotation students from the VAI graduate class of 2025 for helpful discussion and feedback on the manuscript. We thank all members of the Van Andel Institute Genomics Core for technical assistance with library preparation and sequencing. We thank Daniel Bautista and the Van Andel Bioinformatics and Biostatistics Core for help with demultiplexing libraries and establishing the CoCUT&Tag alignment pipeline. We are also grateful for Courtney Zirkle for creative direction on figure panels, and help editing the abstract.

## Author Contributions

D.H.J. and E.B. conceived the study. E.B., H.J.A, A.P.E and D.H.J. performed the experiments; W.S. and D.H.J. performed data analysis. W.S., E.B., and D.H.J. wrote the manuscript; E.B., W.S. and D.H.J reviewed and edited the manuscript, and all authors approved the manuscript.

## Human Samples and ethics statement

Human peripheral blood mononuclear cells were transferred from the Henikoff lab as cryopreserved nuclei stocks, prepared from healthy adult consented donors at the Fred Hutchinson Cancer Center (Institutional Review Board IRB no. 0999.209), as previously described^45^. Human bone marrow mononuclear cells from healthy donors (two males and two females) were obtained from STEMCELL Technologies (cat. no. 20001). All samples were obtained in accordance with the Declaration of Helsinki. Written consent was obtained from all donors to permit the use of their de-identified samples in medical research and for the sharing of primary DNA sequencing data. The study was overseen by the institutional review board of the Van Andel Institute (IR protocol NHS-25005). None of the donors received compensation for their inclusion in this study.

---

**Supplementary Figure 1.**
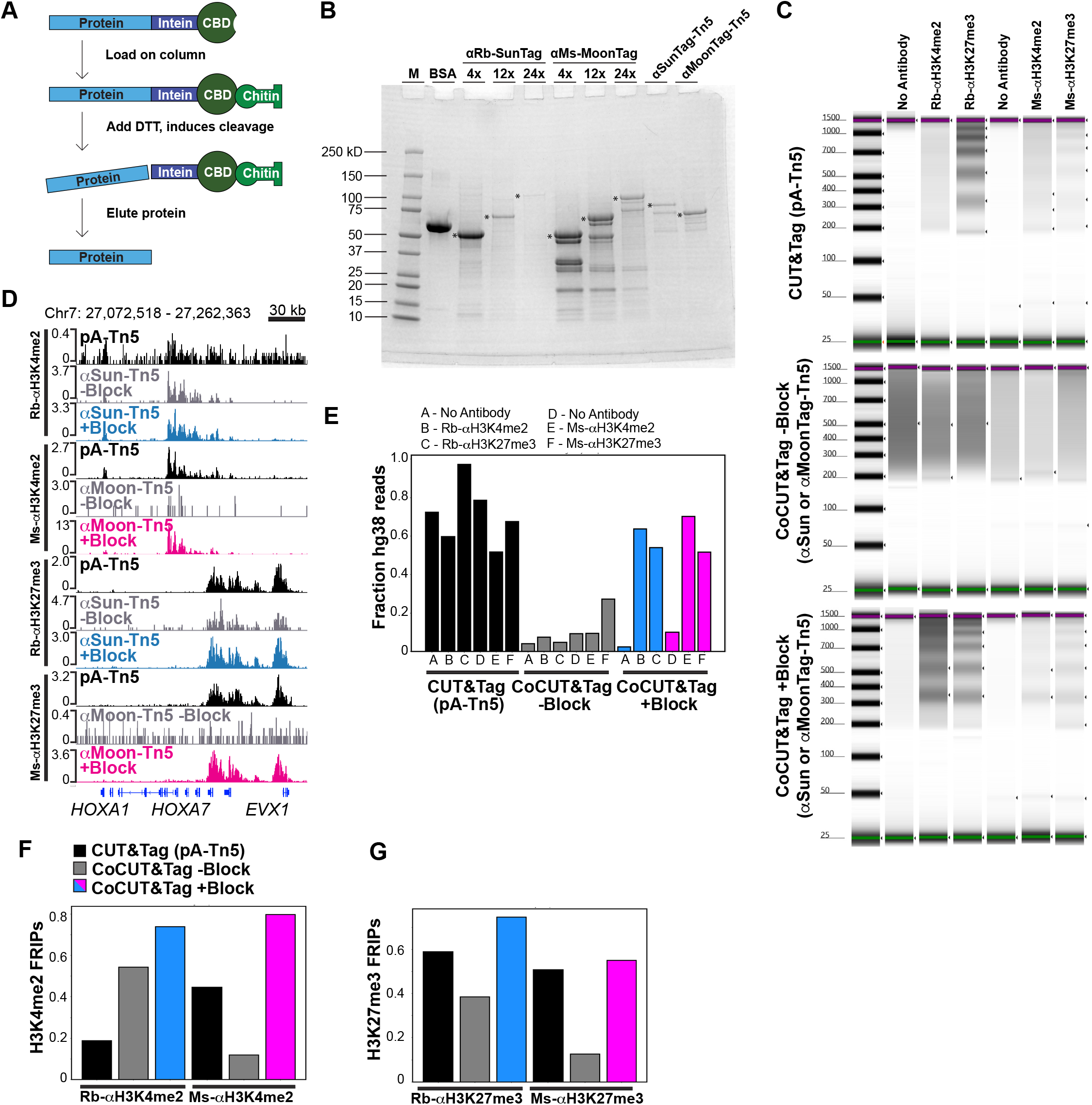
CoCUT&Tag reagent development and optimization for bulk profiling. **(A)** All synthetic proteins in the CoCUT&Tag toolkit were expressed in E.coli fused to an N-terminal intein and chitin-binding domain (CBD), enabling purification by binding to chitin resin followed by DTT induced reduction and cleavage of the intein. **(B)** SDS-Page gel stained with Coomassie showing the 4x, 12x, and 24x anti-rabbit-SunTag and anti-rabbit-MoonTag antigen-peptide repeats and the anti-Sun-Tag-Tn5 and anti-MoonTag-Tn5 fusion proteins. Protein bands of the expected size are indicated by an asterisk. **(C)** Tape station electrophoresis gels showing the fragment-size distributions of Standard CUT&Tag (top) and bulk CoCUT&Tag libraries generated in the absence of blocking agents (-Block; middle) or with 0.5% (w/v) BSA and Casein added to the Wash Buffers (+Block; bottom). The addition of blocking agents restores the nucleosomal laddering pattern in the CoCUT&Tag libraries. **(D)** Genome browser tracks of the *HOXA* locus in RS4;11 cells showing improved recovery of on-target chromatin signal after addition of block. **(E)** Fraction of sequenced reads aligning to the human genome for standard CUT&Tag and CoCUT&Tag libraries generated with or without block. Blocking improves the mapping rate. **(F)** Fraction of reads in peaks (FRiP) for H3K4me2 in blocked and unblocked CoCUT&Tag libraries compared with standard CUT&Tag. After blocking, CoCUT&Tag achieves FRiP values comparable to or better than standard CUT&Tag. **(G)** Same as (F) but showing the FRIPs for H3K27me3.

**Supplementary Figure 2.**
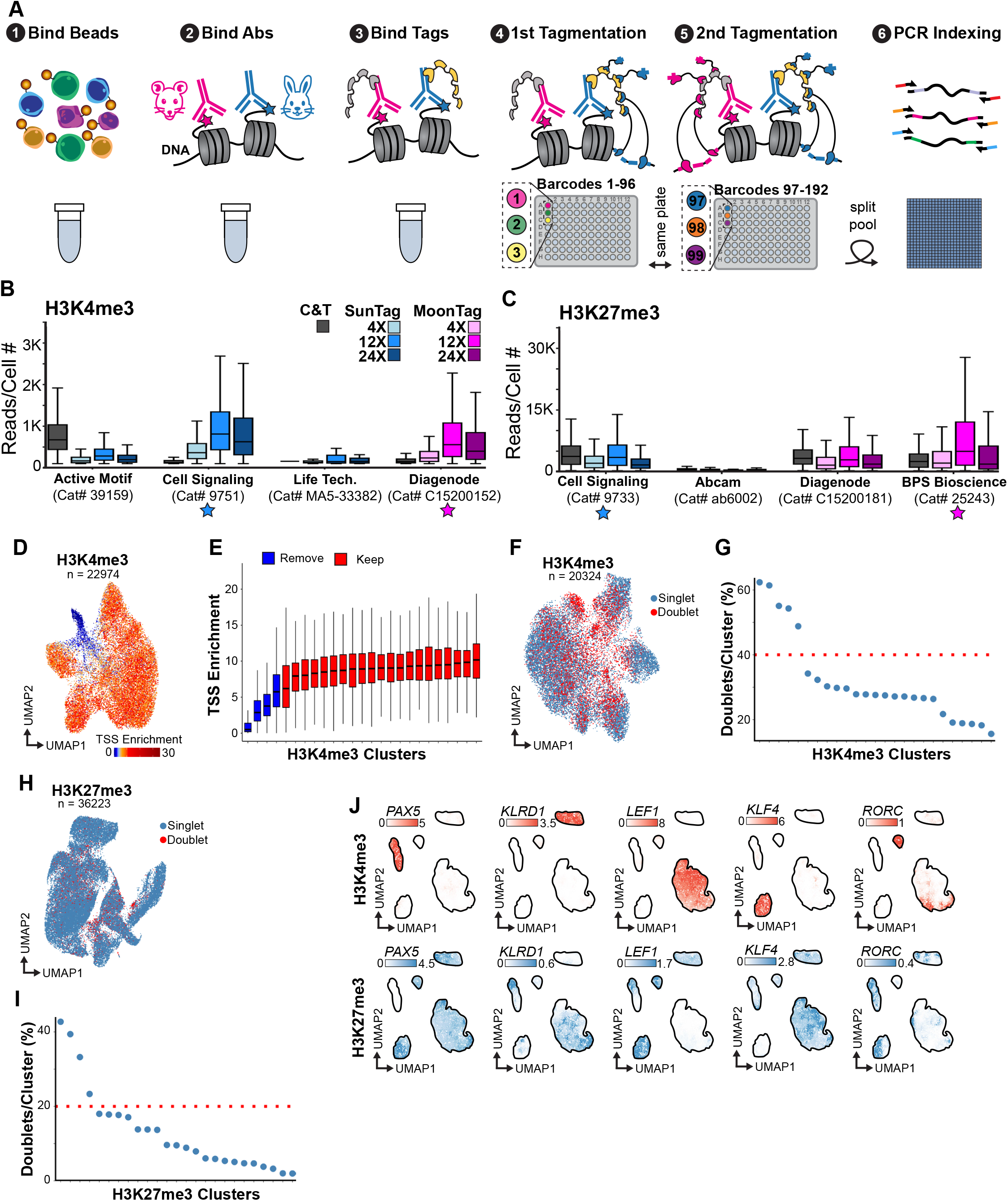
Single-cell CoCUT&Tag workflow, antibody optimization, and PBMC quality control. **(A)** Combinatorial indexing workflow for single-cell CoCUT&Tag. (1) Whole cells were permeabilized, bound to magnetic beads, and lightly crosslinked to prevent aggregation, after which (2) rabbit and mouse primary antibodies were bound in bulk, (3) followed by binding of the corresponding SunTag and MoonTag scaffolds. (4) Cells were then distributed across a 96-well plate for sequential rounds of anti-SunTag-Tn5 binding and tagmentation, (5) followed by anti-MoonTag-Tn5 binding and tagmentation. Keeping cells arrayed in the same plate preserves the cellular linkage between the barcodes from the first and second round of tagmentation. (6) This is followed by split-pool indexing during PCR on a nanowell dispenser. This approach yields up to 50,000 single-cell profiles per reaction, and starting with 1 million cells typically yields enough material for 4 split-pooling reactions. **(B,C)** Antibody and scaffold optimization for H3K4me3 (B) and H3K27me3 (C). Selected rabbit and mouse antibodies perform comparably to or better than standard CUT&Tag when paired with 12x SunTag or MoonTag constructs. **(D,E)** PBMC H3K4me3 quality control. UMAP projection colored by TSS enrichment (D) and cluster-level TSS enrichment used to identify low-quality clusters for removal (E). **(F,G)** Doublet detection and removal for PBMC H3K4me3 data using ArchR-based synthetic doublet scoring and cluster-level doublet enrichment. **(H,I)** Analogous doublet filtering and quality control for PBMC H3K27me3 data. **(J)** PBMC weighted nearest neighbor UMAP-embedding colored by the ArchR-imputed H3K4me3 gene scores (top) and H3K27me3 gene scores (bottom), for marker genes enriched in B-cells (*PAX5*), NK Cells (*KLRD1*), T cells (*LEF1*), monocytes (*KLF4*) and Cytotoxic T cells (*RORC*).

**Supplementary Figure 3.**
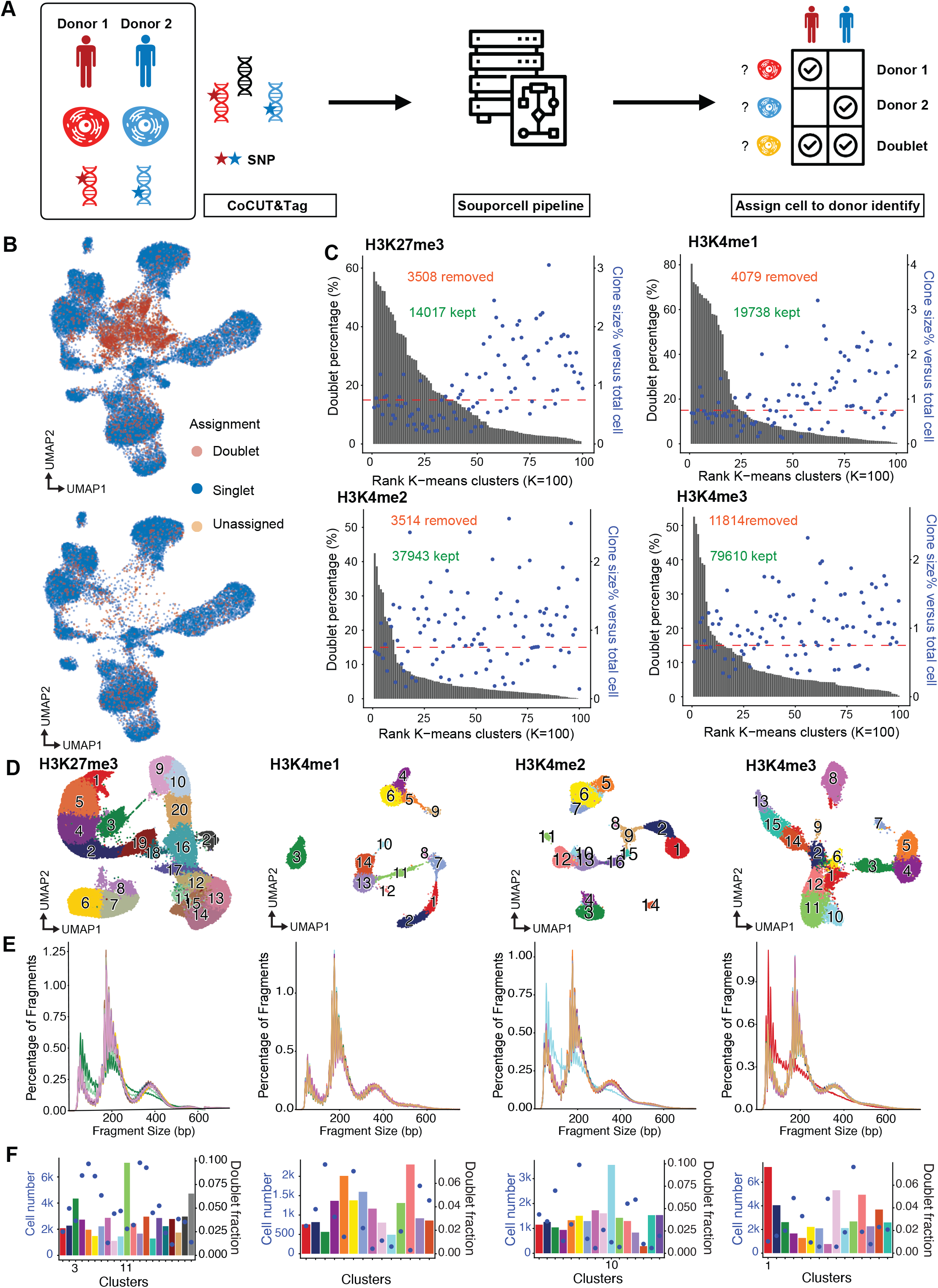
Mixed-donor doublet detection and quality control for bone marrow CoCUT&Tag. **(A)** Experimental and computational workflow for mixed-donor doublet detection. Bone marrow cells from multiple donors were pooled and analyzed using SNP-based donor assignment to identify singlets and doublets. **(B)** UMAP embeddings colored by donor-assigned singlets, doublets, and unassigned cells before and after the first round of cluster-based removal of doublet-enriched populations. **(C)** Quantification of doublet enrichment across K-means clusters for H3K27me3, H3K4me1, H3K4me2, and H3K4me3 datasets, used to identify clusters removed in the first round of filtering. **(D)** Reclustering and embedding after initial doublet-enriched cluster removal. **(E)** Fragment-size distributions of clusters from (D). Residual low-quality clusters show weakened nucleosomal laddering, consistent with degraded chromatin or poor cellular integrity. **(F)** Residual low-quality clusters also retain elevated doublet fractions and were removed in a second round of filtering.

**Supplementary Figure 4.**
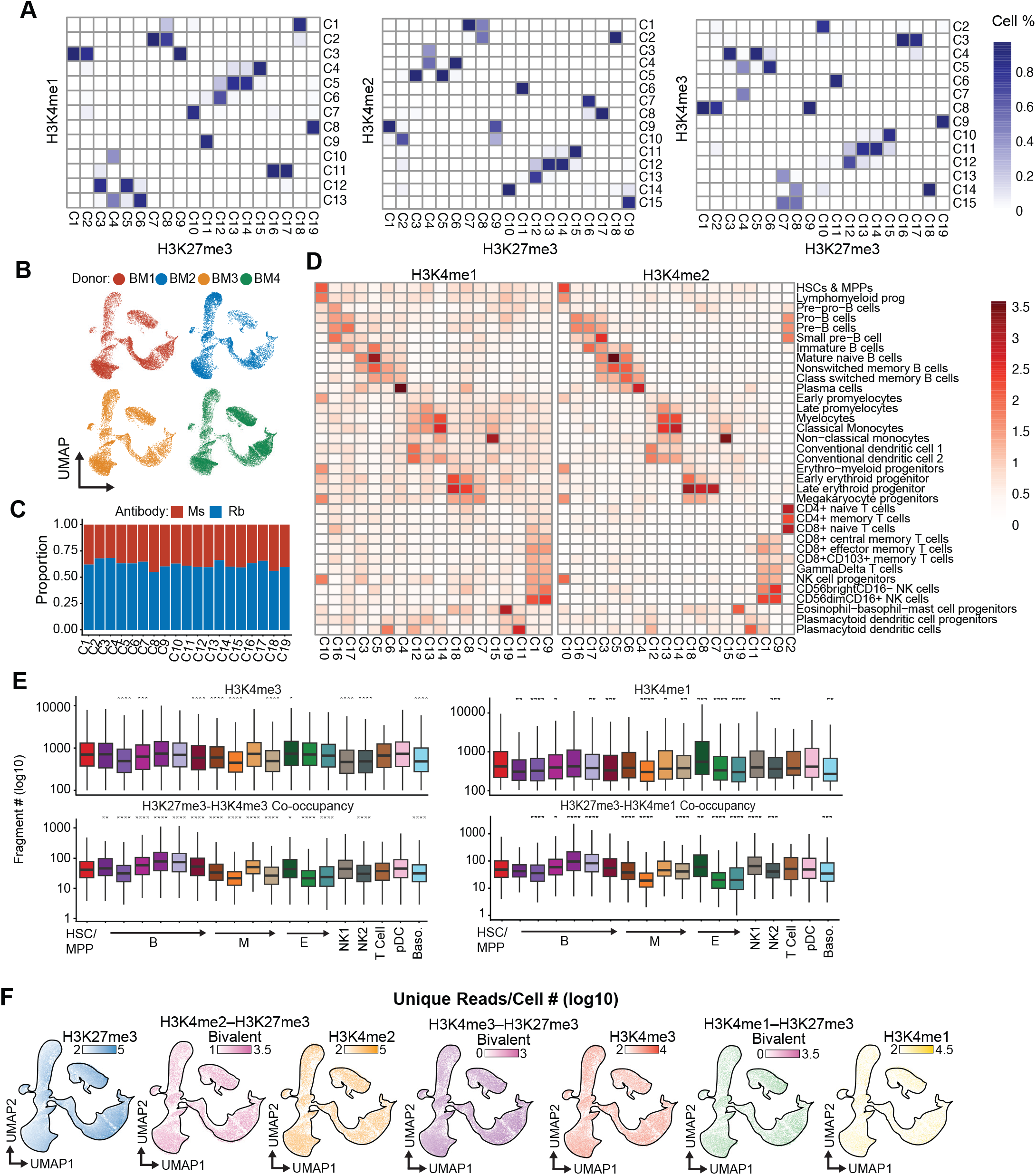
Additional validation of filtered bone marrow CoCUT&Tag datasets. **(A)** Correspondence of cluster identities across paired H3K27me3 and H3K4me1, H3K4me2 and H3K4me3 datasets. Cells from a single cluster in the H3K4 methylation embeddings are often split into multiple clusters by the paired H3K27me3 profiles, reinforcing the added power of H3K27me3 profiles for resolving hematopoietic cell states. **(B)** Distribution of cells from each donor across the H3K27me3 embedding, showing broad donor mixing across hematopoietic populations. **(C)** Distribution of cells profiled with rabbit and mouse H3K27me3 antibodies across clusters, indicating minimal antibody-specific batch effects. **(D)** Heatmaps of cluster-specific ArchR-imputed genes from the H3K4me1 and H3K4me2 datasets alongside expression of the same genes in a reference single-cell RNA-seq dataset with surface marker-defined hematopoietic identities. **(E)** Reads per cell for H3K27me3-H3K4me1 co-occupancy, H3K27me3-H3K4me3 co-occupancy, H3K4me1, and H3K4me3 across retained bone marrow cell states. **(F)** H3K27me3 BMMC UMAP embedding, colored by the unique read counts for each mark.

**Supplementary Figure 5.**
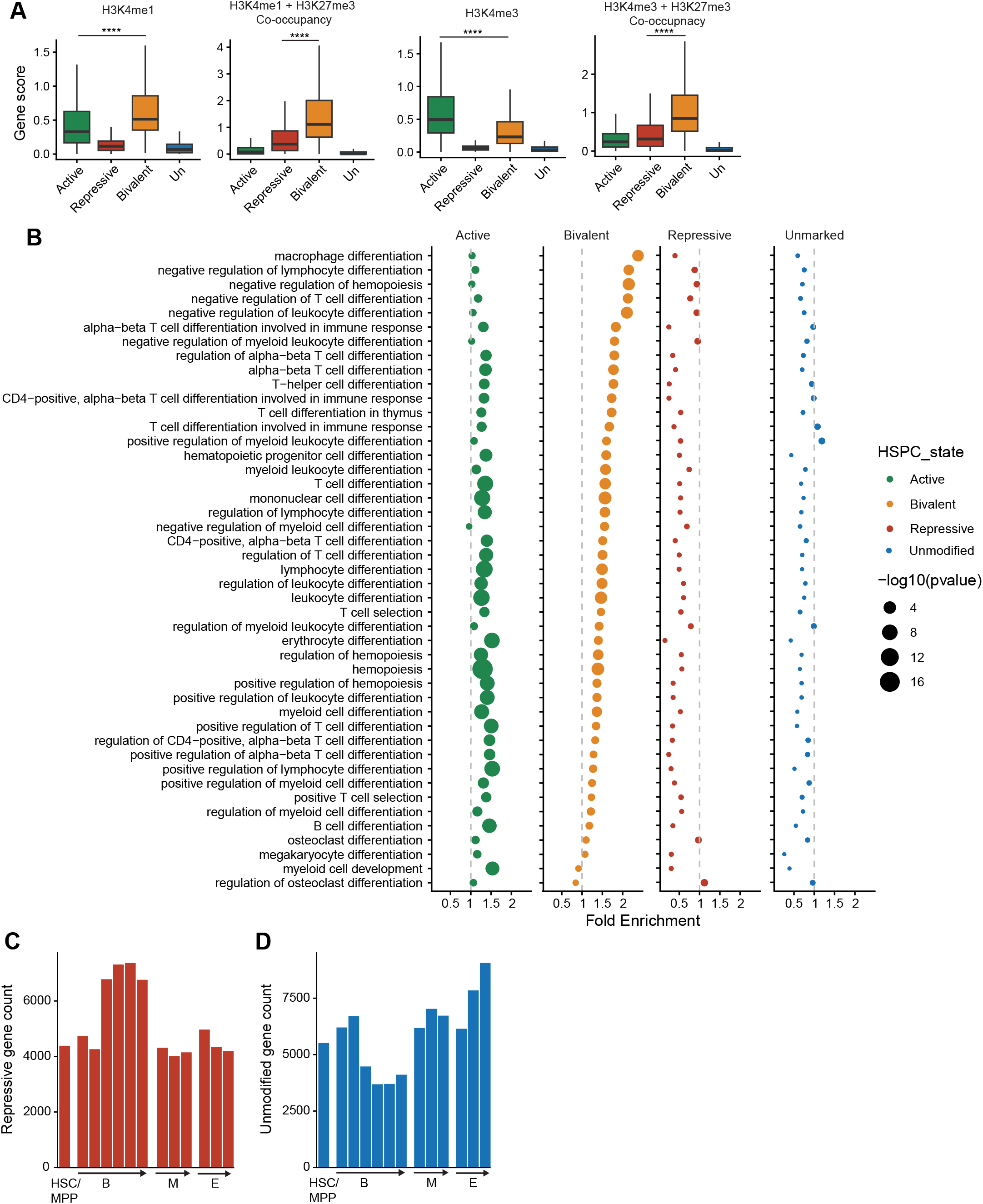
Additional analysis of bivalent, repressed, and unmarked gene states across hematopoiesis. **(A)** Boxplots showing the H3K4me1, H3K4me3 and corresponding H3K27me3 co-occupancy gene scores for genes classified as Active, Repressive, Bivalent and Unmarked. **(B)** Gene Ontology enrichment across active, bivalent, repressed, and unmarked gene classes in HSC/MPPs. Bivalent genes are enriched for hematopoietic developmental programs. **(C)** Absolute number of repressed genes across the HSC/MPP and lineage-committed B, M, and E clusters. **(D)** Absolute number of unmarked genes across the HSC/MPP and lineage-committed B, M, and E clusters.

**Supplementary Figure 6.**
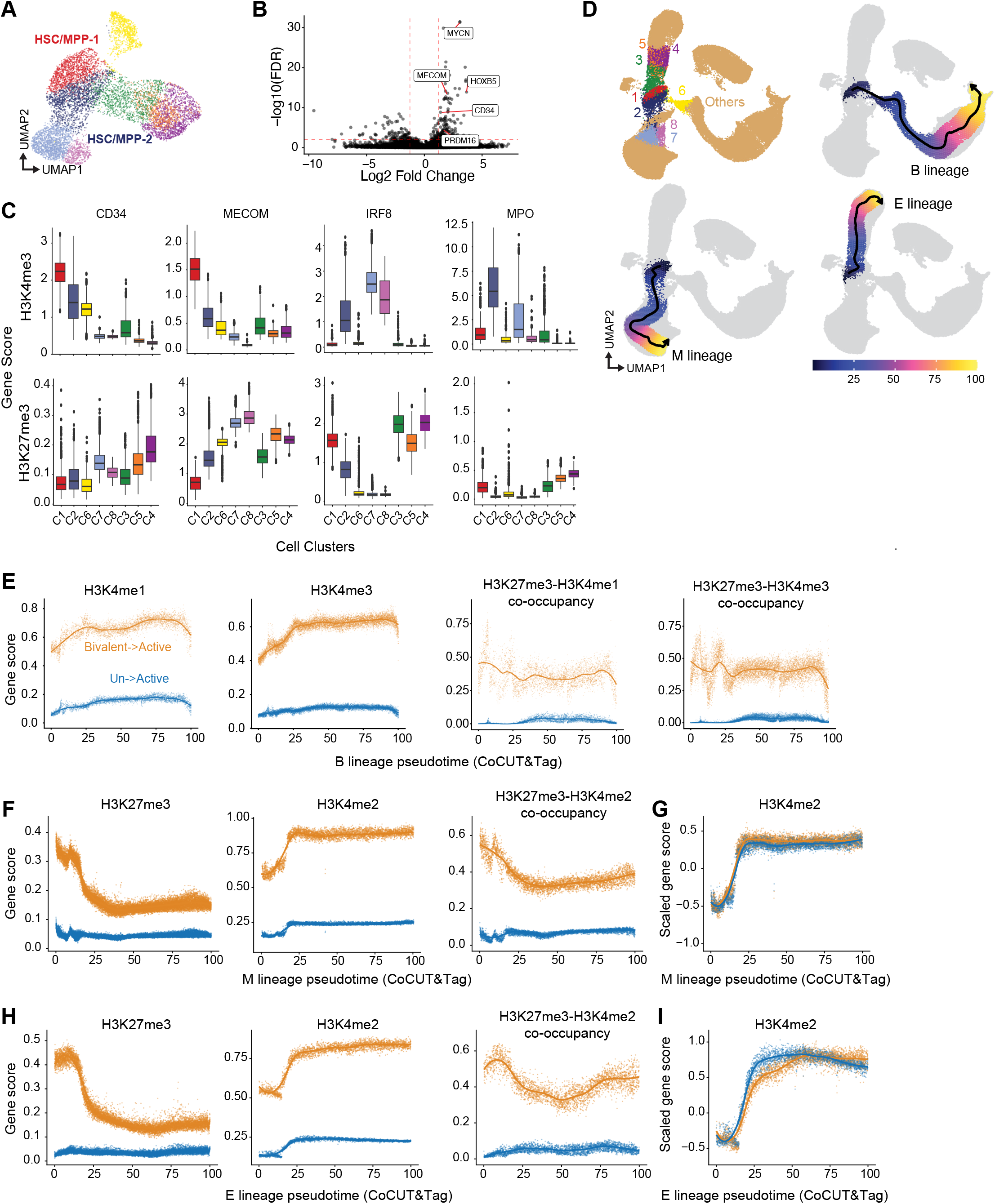
Definition of the HSC/MPP root state and additional pseudotime analyses. **(A)** Refined clustering of the HSC/MPP compartment used to define the root population for developmental trajectory analysis. **(B)** Differentiall analysis of the H3K4me3 gene scores showing the HSC genes *MYCN, MECOM, HOXB5, PRDM16* and *CD34* were enriched for H3K4me3 in the HSC/MPP-1 cluster from (A), and this cluster was defined as the root for trajectory analysis. **(C)** Distribution of representative HSC-associated genes (*MECOM, CD34*) and myeloid-associated genes (*IRF8, MPO*) across the refined HSC/MPP subclusters using H3K4me3 and H3K27me3 ArchR-imputed gene scores. **(D)** B, M, and E lineage pseudotime trajectories overlaid on the integrated hematopoietic embedding. **(E)** Smoothed CoCUT&Tag gene scores for H3K4me1, H3K4me3, H3K27me3-H3K4me1 co-occupancy, and H3K27me3-H3K4me3 co-occupancy across B-lineage pseudotime for genes that transition from bivalent-to-active chromatin or from unmarked-to-active chromatin. **(F)** Same as (E) but showing gene scores across the myeloid-lineage trajectory. **(G)** Scaling the H3K4me2 gene scores from (F) shows that unmarked and bivalent genes gain H3K4me2 with similar timing. **(H)** Same as (E) but showing gene scores across the erythroid-lineage trajectory. **(I)** Same as (G) but showing the scaled H3K4me2 gene scores from the erythroid lineage trajectory.

**Supplementary Figure 7.**
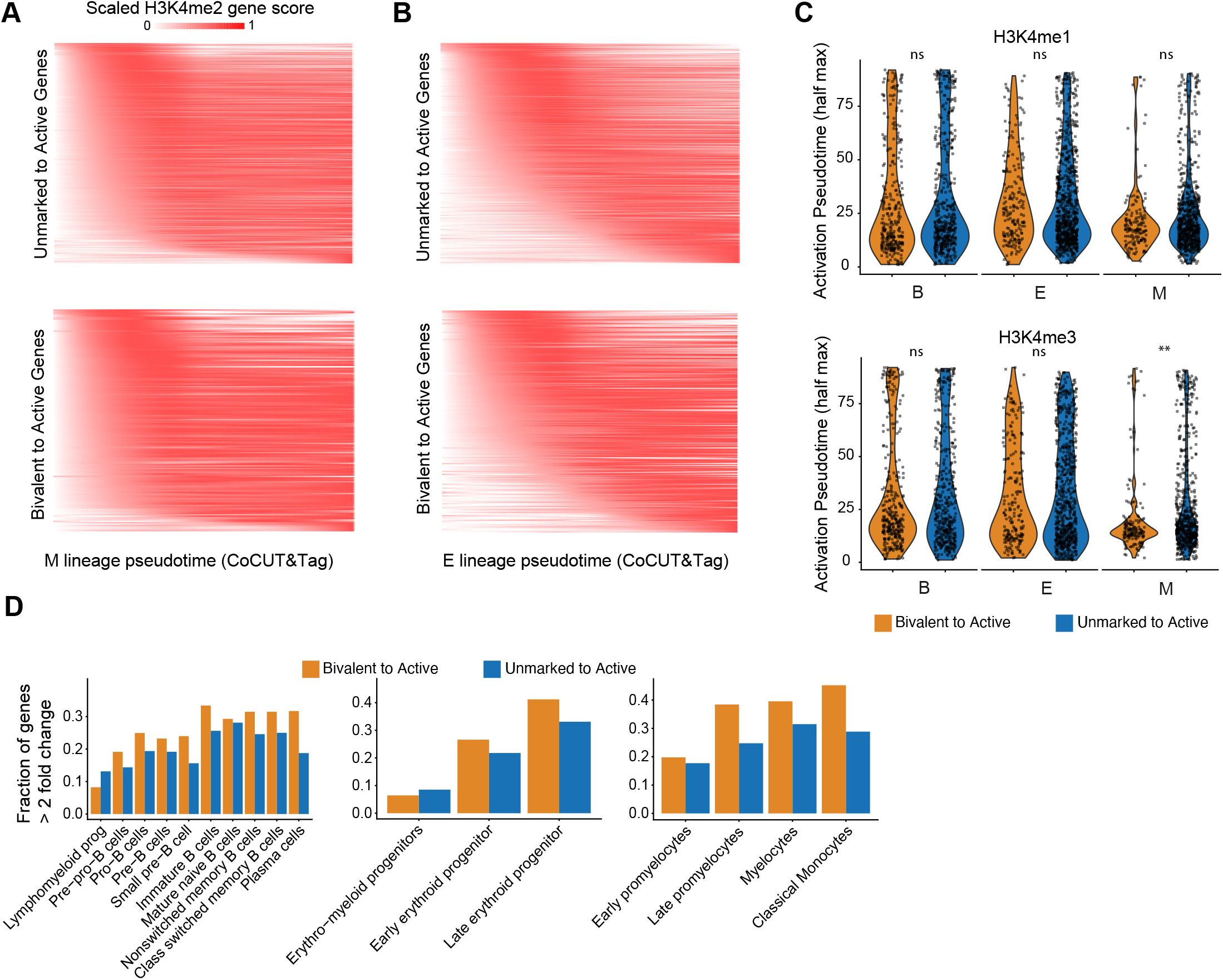
Additional comparisons of gene activation timing and magnitude. **(A)** Heatmap comparing the H3K4me2 gene scores across myeloid-lineage pseudotime for the unmarked genes (top) versus the bivalent genes (bottom) that transitioned to active chromatin. Genes are rank ordered according to the point they reach the half max H3K4me2 gene scores in pseudotime. **(B)** Same as (A) but for the erythroid lineage. **(C)** Violin plot showing the H3K4me1 (top) and H3K4me3 (bottom) activation timing (half-max point in pseudotime) is similar for bivalent and unmarked genes. **(D)** Proportion of genes with greater than two-fold induction across pseudotime in the B, E, and M lineages for genes that transition from bivalent-to-active chromatin or from unmarked-to-active chromatin.

**Supplementary Figure 8.**
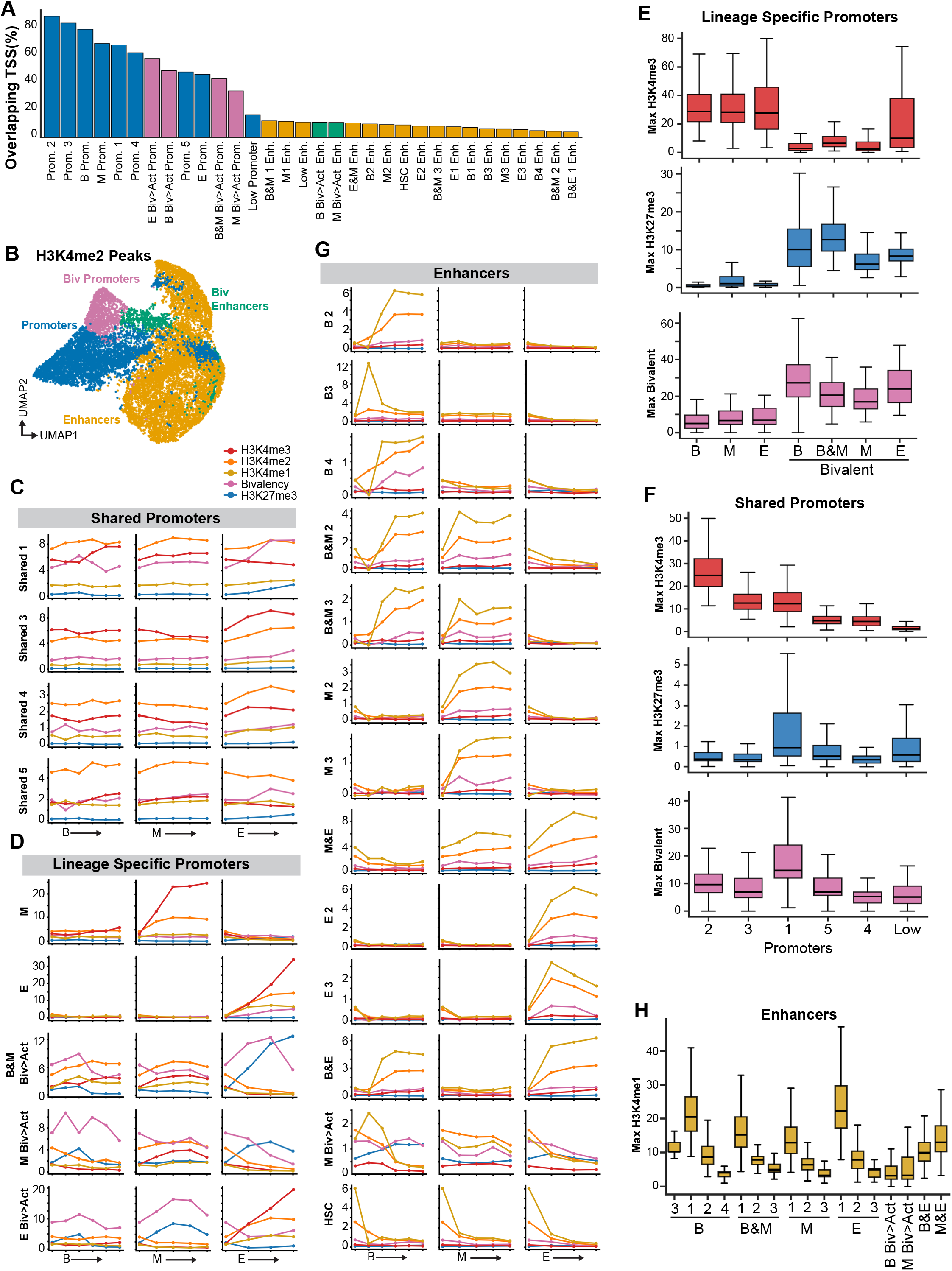
Additional promoter and enhancer subclass definitions. **(A,B)** Additional promoter and enhancer subclass annotations. Promoter classes are strongly enriched near annotated transcription start sites, whereas enhancer classes are predominantly distal. **(C,D)** Average chromatin profiles for general and lineage-specific promoter subclasses across the B, M, and E lineages. Lineage-specific promoters show the highest H3K4me3, whereas bivalent promoters retain elevated H3K27me3 and bivalent signal. **(E,F)** Quantification of H3K4me3, H3K27me3, and bivalent signal across lineage-specific promoter classes (E) and general promoter classes (F). **(G)** Additional putative enhancer subclasses, including lineage-specific enhancer groups with the strongest H3K4me1 signal and rare bivalent enhancer classes. **(H)** Quantification of maximal H3K4me1 signal across enhancer subclasses, showing that the strongest lineage-specific enhancer groups tend to have the highest H3K4me1 levels.

**Supplementary Figure 9.**
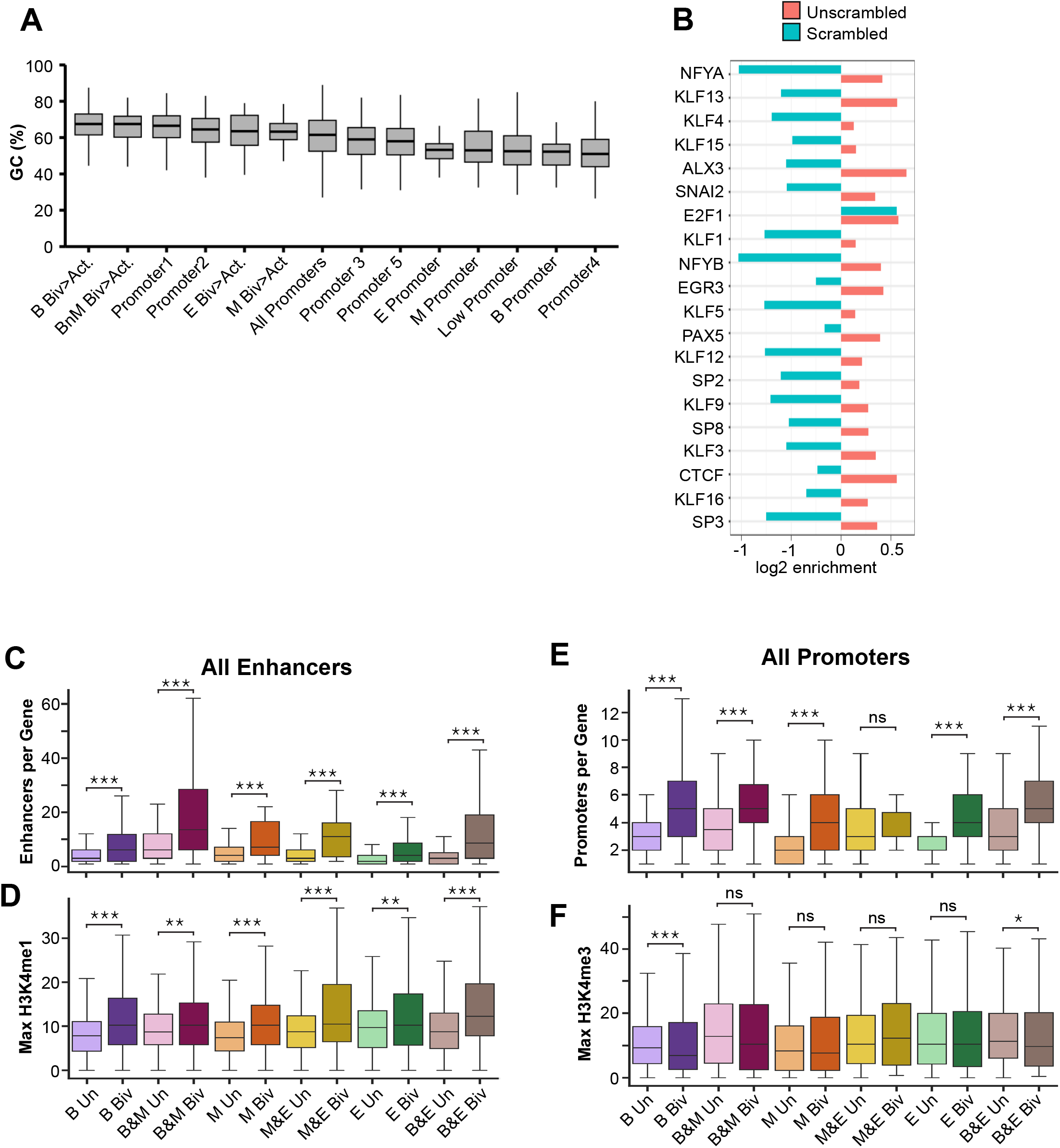
Additional promoter and enhancer features distinguishing bivalent-to-active and unmarked-to-active genes. **(A)** GC content across promoter classes. **(B)** Numerous motifs enriched in the Shared Promoter 1 subclass are GC-rich, including SP/KLF-family factors, CTCF, and PAX5. However, these motifs are no longer enriched after sequence scrambling, indicating their enrichment cannot be explained by GC% alone. **(C,D)** Quantification of enhancer number per gene (C) and enhancer H3K4me1 signal (D) using all enhancer classes rather than only the top lineage-associated enhancer groups. Bivalent-to-active genes retain more enhancers with stronger H3K4me1 signal than unmarked-to-active genes under this broader analysis as well. **(E,F)** Promoter abundance per gene (E) and promoter H3K4me3 signal (F) across gene transition classes. Bivalent-to-active genes tend to reside in more promoter-rich domains than unmarked-to-active genes, but promoters near the bivalent genes are not marked by higher H3K4me3 signal.

**Supplementary Figure 10.**
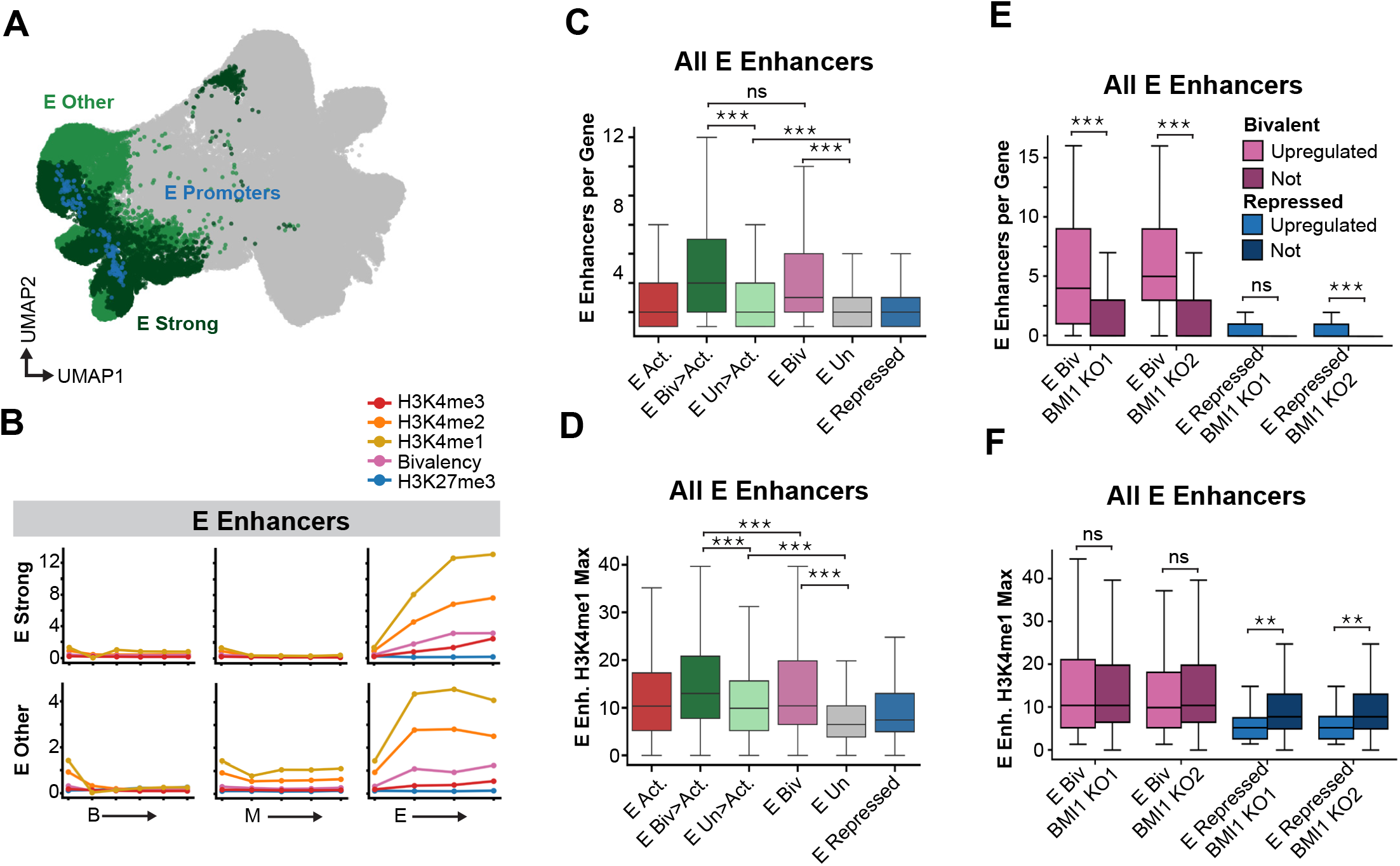
Robustness of erythroid enhancer analyses across enhancer definitions. **(A)** UMAP of all H3K4me3 peaks with strong erythroid enhancers, other erythroid enhancers, and erythroid promoter elements indicated. **(B)** Average chromatin profiles for strong and other erythroid enhancer classes across the B, M, and E lineages. **(C,D)** Number of erythroid enhancers per gene (C) and maximal erythroid enhancer H3K4me1 signal (D) calculated using all erythroid enhancer classes rather than only the strongest enhancer groups. Genes transitioning from bivalent-to-active chromatin retain more erythroid enhancers with stronger H3K4me1 signal than genes remaining bivalent, remaining unmarked, or transitioning from unmarked-to-active chromatin. **(E,F)** *BMI1* knockout analysis repeated using all erythroid enhancer classes. Among genes classified as bivalent in throughout the erythroid lineage, those upregulated after *BMI1* loss carry more erythroid enhancers per gene than non-upregulated bivalent genes, whereas repressed genes do not show the same pattern. Maximal enhancer H3K4me1 signal shows weaker separation than enhancer number.

## Methods

### Cell lines and culture conditions

Human K562 cells (ATCC, Manassas, VA, Catalog #CCL-243) were cultured following the supplier’s protocol in a Sanyo cell culture incubator with standard settings (37°C with 5% CO_2_). RS4;11 cells were a gift from the Bleakley laboratory at the Fred Hutchinson Cancer Center and were transferred from the Henikoff lab as cryopreserved nuclei stocks prepared as previously described^46^.

### CD3 depletion of human bone marrow

CD3-targeted magnetic separation was used to deplete the T cells from bone marrow mononuclear cells (BMMCs). 2.5 × 10^7 BMMCs were thawed in a 37°C water bath and diluted dropwise with 15 mL IMDM (Life Technologies, #12440061) supplemented with 10% FBS (Thermo Fisher Scientific, #A5670801) in a 50 mL conical tube. Cells were passed through a 100 µm MACS SmartStrainer (Miltenyi Biotec, #130-098-463) and centrifuged at 400 × *g* for 5 minutes. The cell pellet was resuspended in 200 µL of wash buffer (1X PBS, 2 mM EDTA, 0.5% BSA) and transferred to a 1.5 mL tube. Mixed CD3 MicroBeads (50 µL; Miltenyi Biotec, #130-050-101) were added, and samples were incubated at 4°C for 15 minutes. Following incubation, samples were washed with 1.25 mL wash buffer, centrifuged at 300 × *g* for 5 minutes, and resuspended in 500 µL wash buffer.

For magnetic separation, an LS column (Miltenyi Biotec, #130-042-401) was equilibrated with 3 mL wash buffer. Samples were applied to the column, and the CD3-negative fraction (T cell-depleted BMMCs) was collected in a 15 mL conical tube. The column was rinsed twice with wash buffer, and the flow-through was collected in the same tube. Cell viability and concentration were determined using a Countess cell counter. Cells were cryopreserved at 1e6 cells/mL in freezing medium (45% FBS, 45% IMDM, 10% DMSO), by placing them in a Mr. Frosty isopropanol chamber and transferring to a -80°C freezer. For long-term storage, cells were transferred to a liquid nitrogen tank.

### Antibodies

Of the rabbit and mouse monoclonal antibodies we tested, we found the following antibodies performed the best in CoCUT&Tag assays: rabbit anti-H3K4me3 (Cell Signaling Technologies; cat. no. 9751), mouse anti-H3K4me3 (Diagenode; cat no. C15200152), rabbit anti-H3K27me3 (Cell Signaling Technologies; cat. no. 9733), mouse anti-H3K27me3 (BPS Bioscience; cat. no. 25243), rabbit anti-H3K4me2 (Cell Signaling Technologies; cat. no. 9725), mouse anti-H3K4me2 (Diagenode; cat. no. C15200151), rabbit anti-H3K4me1 (Cell Signaling Technologies; cat. no. 5326), mouse anti-H3K27ac (Diagenode; cat. no. C15200184). In addition, we performed standard CUT&Tag assays and CoCUT&Tag assays with the following antibodies: rabbit anti-H3K4me3 (Active Motif; cat. no. 39159), mouse anti-H3K4me3 (Life Technologies; cat. no. MA5-33382), mouse anti-H3K27me3 (Abcam; cat. no. ab6002), mouse anti-H3K27me3 (Diagenode; cat. no. C15200181), rabbit anti-H3K4me2 (Sigma Aldrich; cat. no. 07-030), mouse anti-H3K4me2 (Active Motif; cat. no. 39679), mouse anti-H3K4me1 (Life Technologies; cat. no. MA5-23514). For standard CUT&Tag assays using a rabbit primary antibody, the secondary guinea pig anti-rabbit IgG antibody (Antibodies Online, cat. no. ABIN101961) served as an adaptor to amplify the number of tethered pA-Tn5 complexes. To amplify the signal from mouse primary antibodies, we used the secondary rabbit anti-mouse IgG antibody (Abcam; cat. no. ab46540). In CoCUT&Tag assays, the anti-rabbit-IgG-SunTag, and anti-mouse-IgG-MoonTag scaffold proteins replace the secondary antibodies for signal amplification.

### Cloning plasmids for bacterial expression of CoCUT&Tag proteins

The backbone plasmid used for all cloning was the 3XFlag-pA-Tn5-Fl vector (Addgene #124601), which was gifted to us by Steve Henikoff’s lab. To enable flexible targeting of the SunTag and MoonTag to chromatin proteins, we fused the SunTag and MoonTag to anti-Rabbit-IgG (TP897) and anti-Mouse-IgG (TP1170) nanobodies^18^. We ordered the anti-rabbit::4xSunTag::sfGFP::GB1 gene and the anti-mouse::4xMoonTag::mCherry2 gene as synthesized gBlock HiFi Gene Fragments from IDT. The GB1 domain was later removed from the anti-Rabbit-SunTag-sfGFP construct because it causes unwanted binding to Mouse IgG. We used Gibson Assembly to insert each gene into the backbone plasmid, completely replacing the 3XFlag-pA-Tn5-Fl protein. We then ordered an 8x and 20x repeat of both the SunTag and MoonTag sequences from GenScript and used restriction cloning to add the additional repeats into the 4xSunTag and 4xMoonTag constructs. This strategy generated a 4x, 12x, and 24x antigen-peptide repeat version of both the anti-Rabbit-SunTag and anti-Mouse-MoonTag constructs. We found the 12x constructs are ideal for CoCUT&Tag assays and we deposited the corresponding vectors in Addgene: pTBX1-αRabbit-12xSunTag-sfGFP (Addgene plasmid # 254519) and pTBX1-αMouse-12xMoonTag-mCherry2 (Addgene plasmid # 254520). To generate the complimentary Tn5 fusion proteins, we ordered the anti-SunTag and anti-MoonTag genes as synthesized gBlocks Gene Fragments from IDT. We used Gibson assembly to replace the 3xFlag-pA section of the backbone plasmid. We also deposited the completed pTBX1-αSunTag-Tn5 (Addgene plasmid # 254521) and pTBX1-αMoonTag-Tn5 (Addgene plasmid # 254522) constructs into Addgene.

### Protein purification

All the synthetic proteins for CoCUT&Tag were expressed with a C-terminus fusion to an intein and chitin binding domain (CBD). The CBD specifically binds chitin resin, and addition of a reducing agent (Dithiothreitol, DTT) causes the intein to cleave and release the target protein into solution. All the CoCUT&Tag proteins were obtained using the same protein expression and purification workflow.

Each construct was transformed into T7 Express lysY/Iq Competent *E. coli* (NEB #C3013I) following the manufacturer’s protocol and plated for selective overnight growth. A single colony was used to inoculate 5 mL of LB media containing 100 µg/ml carbenicillin. The starter culture was incubated overnight at a temperature of 37°C while shaking at 220 RPM. The 5 mL starter culture was then used to inoculate 345 mL TB media containing 100 µg/ml carbenicillin. The 350 mL culture was incubated at a temperature of 37°C while shaking at 170 RPM until the OD600 reached 0.75-1.00 (approximately two hours). After reaching this density, the culture was placed in a cold room for one hour to cool then promptly induced via the addition of IPTG to a final concentration of 0.25 mM. The induced culture was incubated overnight at a temperature of 18°C while shaking at 170 RPM. The bacterial culture was pelleted by centrifugation for 10 minutes at 9,500 × *g*. All centrifugations were performed at a temperature of 4°C. The bacterial pellet was flash frozen using liquid nitrogen and stored at a temperature of -80°C. To proceed, the pellet was thawed on ice in a 4°C cold room then resuspended in 35 mL chilled HEGX (0.02 M HEPES-KOH at pH 7.2, 1 M NaCl, 0.001 M EDTA, 10% glycerol, 0.2% Triton X-100) with fresh protease inhibitor added immediately before use (Sigma #5056489001) using a stir plate. The resuspension was sonicated in ice-nested beakers with a microtip set to 40% amplitude for 12 rounds: 3 seconds on, 3 seconds off, for 40 seconds total as detailed by Bryson & Henikoff (https://dx.doi.org/10.17504/protocols.io.8yrhxv6). The lysate was then centrifuged at 29,000 × *g* for 30 minutes. A chitin resin column was prepared in a 4°C cold room by gently inverting the resin (NEB #S6651) to resuspend and adding 2.5 mL resin to an Econo-Pac Chromatography Column (Bio-Rad #7321010). With the column set upright in a column rack, the resin was allowed to settle and then washed twice with 20 mL HEGX Buffer. Once the HEGX Buffer had run through the column, the bottom of the column was capped, and 20 mL of the soluble lysate (i.e., supernatant) was added. The column was capped on the top, parafilmed, and left nutating overnight at 4°C. The next day, the column was returned to the rack, and the unbound fraction was drained into a collection tube. After washing the column twice with 20 mL HEGX Buffer and recapping the bottom, 6 mL HEGX Buffer containing 100 mM DTT and fresh protease inhibitor was added slowly to the column. The column was capped on the top, parafilmed, and incubated for 48 hours at 4°C. The DTT-cleaved fraction was then drained into a collection tube. The collected sample was loaded in a dialysis cassette (Life Tech #66810) following product literature and dialyzed overnight in Dialysis Buffer (0.1M HEPES-KOH at pH 7.2, 0.2 M NaCl, 0.2 mM EDTA, 0.2% Triton X-100, 20% glycerol, with DTT added to a final concentration of 1.7 mM immediately before use) with a buffer change after two hours. The dialyzed sample was recovered from the cassette and centrifuged at 3000 × *g* for two minutes to remove precipitated protein. The sample was then held on ice at 4°C until the protein was verified on an SDS-PAGE gel. After verification, the Tn5-fusion proteins were loaded with adapter oligos and stored, as is further described. The solution containing Tn5-fusion proteins can also be snap frozen and stored at -80°C for loading up to one year later. For all other proteins, glycerol was added to 50% and the protein was stored at -20°C.

### Anti-SunTag-Tn5 and Anti-MoonTag-Tn5 adapter loading

Prior to starting the experiment, the anti-SunTag-Tn5 and anti-MoonTag-Tn5 were loaded with differentially barcoded ME-s5 and ME-s7 adapters, using a similar protocol and the same adapter sequences to the one described previously for sciCUT&Tag and sciCUT&Tag2in1^3,45^. These barcodes are essential to distinguish genomic reads corresponding to protein 1 from genomic reads corresponding to protein 2 and the co-occupancy reads.

Briefly, the differentially barcoded ME-s5 and ME-s7 and the universal ME-reverse oligo were ordered from IDT (PAGE purification is used for the ME-s5 and ME-s7 oligos). For bulk CoCUT&Tag, we used 4 ME-s5 oligos and 4 ME-s7 oligos to load anti-SunTag-Tn5 and a different combination of 4 ME-s5 oligos and 4 ME-s7 oligos to load anti-MoonTag-Tn5. This ensures sufficient adapter diversity for sequencing on Illumina or Element Aviti instruments.

For single-cell combinatorial indexing CoCUT&Tag workflows, we prepared a 96 well plate of differentially barcoded anti-SunTag-Tn5s and a second 96 well plate of differentially barcoded anti-MoonTag-Tn5s. The anti-SunTag-Tn5 plate was loaded with an 8X12 grid of ME-s5 oligos with barcodes 1-8, and ME-s7 oligos with barcodes 1-12. The anti-MoonTag-Tn5 plate was loaded with an 8X12 grid of ME-s5 oligos with barcodes 9-16, and ME-s7 oligos with barcodes 13-24. Briefly, the lyophilized oligos are resuspended to a final concentration of 200 μM in annealing buffer (10mM Tris pH8, 50mM NaCl, 1 mM EDTA). We then mixed 200 μM ME-s5 oligo with 200 μM ME-reverse oligo resulting in 100 μM ME-s5 adapter, and mixed 200 μM ME-s7 oligo with 200 μM ME-reverse resulting in 100 μM ME-s7 adapter.

To anneal the adapters, we placed the tubes in a thermocycler and ran an annealing program with a 105°C heated lid that started with a 95°C 2-minute incubation and decreases the temperature in 5°C increments incubating 5 minutes each, ending with a 5 minute incubation at 25°C and the hold at 8°C. Annealed oligos were kept on ice for immediate use or stored at −20°C. Annealed adapter stocks can be kept at −20°C for at least a year. The annealed ME-s5 and ME-s7 adapters were then mixed at equimolar concentration and suspended in anti-SunTag-Tn5 or anti-MoonTag-Tn5 solution at a ratio of 1 ME-s5:1 ME-s7:12.5 Tn5-fusion protein by volume. This ensures the adapters are in excess. The efficiency of the CoCUT&Tag reactions using these reagents is dependent on the concentration of the anti-SunTag-Tn5 and anti-MoonTag-Tn5 proteins, with high concentrations of proteins translating into higher yields until saturation is reached. For example, for loading the 96 well anti-SunTag-Tn5 plate, we used a multichannel pipette to distribute 1.2 μL of ME-s5 1-8 across rows A-H of a 96 well plate. We then distributed 1.2 μL of ME-s7 1-12 across columns 1-12 of the 96 well plate. We then distributed the anti-SunTag-Tn5 protein prep into a reservoir and used a multichannel pipette to add 15 μL to each well of the 96 well plate, pipetting up and down 3 times after dispensing the protein to ensure rapid mixing and even adapter loading. We then sealed the plate. After combining the annealed adapters and Tn5-fusion proteins, the solution was vortexed to mix, and loading was allowed to proceed for 1.5 hours at room temperature. An equal volume of 90% glycerol was then added to each tube or well and the loaded anti-SunTag-Tn5 and anti-MoonTag-Tn5 solutions were stored at -20°C.

### Bulk CoCUT&Tag Protocol

Prior to starting the protocol, we prepared 50 g/L Casein Block Buffer, pH 7.0 by adding 5.99 g NaOH to 400 mL of nanopure water. We then slowly added 25 grams of Casein while stirring to avoid clumping. This solution was incubated overnight at 37°C while shaking at 250 RPM to ensure that the protein fully dissolves. We then adjusted the solution to pH to 7.0 by adding 12 N HCl dropwise with a pH meter, with heavy stirring to avoid precipitation. This solution was adjusted to 500 mL aliquoted into 15 mL conical tubes and flash freeze in liquid nitrogen and store at -80°C.

Bangs ConA beads (Bangs Cat. No. BP531) were prepared for binding (5 μL per CoCUT&Tag sample) by transferring them to a 1.5 mL microfuge tube and adding 500 µL of Binding Buffer (400 μL of 1 M HEPES-KOH pH 7.9 [20 mM], 200 μL of 1 M KCl [10 mM] 20 μL of 1 M CaCl2 [1 mM], and 20 μL of 1 M MnCl2 [1 mM], adjusted to 20 mL nanopure H2O). The mixture was added to a magnet stand and the supernatant removed. The beads were washed with another 500 μL of binding buffer and then resuspended in 100 μL of binding buffer.

Whole cells were thawed at room temperature and mixed. For each sample, 25K cells were added to a 1.5 mL tube and centrifuged at 600 × *g* for 3 minutes. The supernatant was removed and the pellet was resuspended in 900 μL of Wash Buffer (48 mL ddH_2_O, 1 mL 1M HEPES pH 7.5[20 mM], 1.5 mL 5M NaCl [150 mM], 12.5 µL 2M Spermidine [0.5 mM], 250 μL of 10% Triton X-100 [0.05%],1 Roche Complete EDTA-free protease tablet).

The binding buffer containing Bangs ConA beads was added to the whole cell suspension in 3 increments of 33 μL, with mixing in between each addition. After 10 minutes of binding at room temperature, the sample was split into 500 μL low bind tubes, 1 tube for each sample, and the supernatant was removed and the suspension was brought up in 15 μL of Antibody Buffer (1 mL Wash Buffer, 4 μL 0.5M EDTA [2 mM]). 1.5 μL of appropriate mouse antibody and 1.5 μL of the appropriate rabbit antibody was added for overnight incubation at 4°C.

Before starting the second day we prepared Wash Block Buffer (48 mL ddH_2_O, 1 mL 1M HEPES pH 7.5[20 mM], 1.5 mL 5M NaCl [150 mM], 12.5 µL 2M Spermidine [0.5 mM], 250 μL of 10% Triton X-100 [0.05%],0.25 g BSA (Sigma-Aldrich Cat no. A9647), 5 mL 50 g/L mL Casein Block Buffer pH 7.0, 1 Roche Complete EDTA-free protease tablet), 300-Wash Block Buffer (48.5 mL Wash Block Buffer, 1.5 mL 5M NaCl [300 mM NaCl]) and TAPS buffer (500 μL 1M TAPS pH 8.5 [10 mM] in 49.5 mL ddH_2_O), and Tagmentation Buffer (9.9 mL 300-Wash Block Buffer, 100 μL 1 M MgCl2 [10 mM]).

Samples were washed twice in 100 μL of Wash buffer and suspended in 50 μL of Wash Block Buffer for 30 minutes at room temperature. In addition, we pre-blocked the SunTag and MoonTag scaffold reagents. For each sample, we mixed 20 μL of Wash Block Buffer with 2 μL of αRb-12X-SunTag-GFP and 2 μL of αMs-12X-MoonTag-GFP and incubated at room temperature for 30 minutes. The supernatant was removed from the cells, and 20 μL of the Block Tag mix was added to the cells for incubation at 4°C for 1 hour.

Samples were washed twice in 100 μL of Wash Block Buffer and brought up in 15 μL of 300-Wash Block Buffer. 1.5 μL aSun-Tn5 loaded with barcoded ME-s5s and ME-s7s were added for incubation at 4°C for 1 hour. Samples were washed twice in 100 μL of 300-Wash Block Buffer and brought up in 50 μL of Tagmentation buffer. The samples were placed in a thermocycler with a heated lid and tagmentation was performed at 37°C for 1 hour.

Samples were washed twice in 100 μL of 300-Wash Block Buffer and brought up in 15 μL of 300-Wash Block Buffer. 1.5 μL aMoon-Tn5 loaded with barcoded ME-s5s and ME-s7s was added for incubation at 4°C for 1 hour. To distinguish between targets and co-occupancy reads, the aSun-Tn5 and aMoon-Tn5s must be loaded with distinct combinations of ME-s5s and ME-s7s adapters.

Samples were washed twice in 100 μL of 300-Wash Block Buffer and brought up in 50 μL of Tagmentation buffer. The samples were placed in a thermocycler with a heated lid and tagmentation was performed at 37°C for 1 hour.

0.1% SDS Release Buffer was prepared (220 μL 10 mM TAPS, 25 μL 1% SDS, 5 μL Thermolabile Proteinase K). Samples were then washed once with 200 μL of 10 mM TAPS pH 8.5. This TAPS wash is to remove salt to prepare the sample for PCR, and so we made sure no excess liquid remained in the cap of the tube or on the walls. The supernatant was then removed and 5 μL of 0.1% SDS Release Buffer was added to each sample, ensuring that the beads are suspended in this small volume by flicking the tube or vortexing. The chromatin release step was performed by incubation at 1 hour at 37°C and 1 hour at 58°C in a thermocycler with a heated lid.

Samples were quenched by addition of 4 μL of 1.5% TritonX-10 solution. This was followed by addition of 24 µL of nanopore H_2_0 to each sample, followed by 18 µL of PCR “Master Mix” (for each sample: 10 µL 5 X HiFi buffer, 5.5 µL ddH_2_O, 1.5 µL 10 mM dNTP, 1 µL KAPA HiFi Polymerase). We then added 2 µL of a barcoded Forward primer (10 µM) and 2 µL of a barcoded Reverse primer (10 µM) to each sample, ensuring that each sample had at least one distinctly barcoded primer. Mix by vortexing. PCR was conducted on a thermocycler with a heated lid using the profile: 58°C for 5 min (gap fill), 72°C for 5 min, 98°C for 45 seconds, followed by 14 cycles of (98°C for 15 seconds, 60°C for 10 seconds, 72°C for 1 seconds), ending with 72°C for 1 minute and hold at 12°C. Samples are then purified using a standard Ampure bead cleanup with a 1.3 X size selection eluted in 30 μL Tris-HCl pH 8.0 and then run on a Tapestation and pooled at equimolar concentrations. The library pool was then subjected to an Ampure bead cleanup with a 0.9 X size selection to remove the PCR product of “internal tagmentation” that results from one adapter tagmenting into another. Sequencing is performed on an Element Aviti or Illumina Platform using the appropriate custom primers.

### Single-cell CoCUT&Tag

We typically start the single-cell CoCUT&Tag protocol with 1-2 million cells, but the protocol can be run with fewer starting cells, the risk being the nanowell chip may need to be loaded with fewer than 10 cells per well. Regardless of the starting cell number, Magnefy ConA beads (Bangs Cat. No. MFY531) were prepared for binding by mixing the solution and adding 200 µL to 500 µL of Binding Buffer (400 μL of 1 M HEPES-KOH pH 7.9 [20 mM], 200 μL of 1 M KCl [10 mM] 20 μL of 1 M CaCl2 [1 mM], and 20 μL of 1 M MnCl2 [1 mM], adjusted to 20 mL nanopure H_2_O). The mixture was added to a magnet stand and the supernatant removed. The beads were washed with another 500 µL of binding buffer and then resuspended in 100 μL of binding buffer.

Whole cells were thawed at room temperature and mixed to a final count of 1-2 million. Cells were added to a 1.5 mL tube and centrifuged at 600 × *g* for 3 minutes. The supernatant was removed and the pellet was resuspended in 900 mL of Wash Buffer (48 mL ddH_2_O, 1 mL 1M HEPES pH 7.5 [20 mM], 1.5 mL 5M NaCl [150 mM], 12.5 µL 2M Spermidine [0.5 mM], 250 μL of 10% Triton X-100 [0.05%],1 Roche Complete EDTA-free protease tablet).

The binding buffer was added to the whole cell suspension in 3 increments of 33 μL, with mixing in between each addition. After 10 minutes of binding at room temperature, the supernatant was removed and the suspension was brought up in 500 μL of Wash buffer. The cells were lightly crosslinked by adding 3.2 μL of 16% Formaldehyde (Thermo Scientific Cat no. 28906) to a final concentration of 0.1%. After 2 minutes, the reaction was quenched by an addition of 15 μL of 2.5 M glycine. To ensure cross-linking was stopped, the samples were then washed immediately with 200 μL of Wash buffer and brought up in 100 μL of Antibody Buffer (1 mL Wash Buffer, 4 μL 0.5M EDTA [2 mM]). At this point the sample could be split for multiple antibody combinations, and the fraction of cells split into each sample should be proportional to the fraction of the 96-well plate that will be dedicated to that sample. Antibody is added at a 1:10 dilution, and in this example since the experiment was run on a single 100 μL sample, 10 μL of the appropriate mouse antibody and 10 μL of the appropriate rabbit antibody was added for overnight incubation at 4°C.

Wash Block Buffer (48 mL ddH_2_O, 1 mL 1M HEPES pH 7.5 [20 mM], 1.5 mL 5M NaCl [150 mM], 12.5 µL 2M Spermidine [0.5 mM], 250 μL of 10% Triton X-100 [0.05%],0.25 g BSA (Sigma-Aldrich Cat no. A9647), 5 mL 50 g/L mL Casein Block Buffer pH 7.0 (preparation described in the Bulk CoCUT&Tag section), 1 Roche Complete EDTA-free protease tablet), 300-Wash Block Buffer (48.5 mL Wash Block Buffer, 1.5 mL 5M NaCl [300 mM NaCl]) and TAPS buffer (500 μL 1M TAPS pH 8.5 [10 mM] in 49.5 mL ddH_2_O), and Tagmentation Buffer (9.9 mL 300-Wash Block Buffer, 100 μL 1 M MgCl2 [10 mM]) were prepared.

Samples were washed twice in 100 μL of Wash buffer and suspended in 50 μL of Wash Block Buffer for 30 minutes at room temperature. 120 μL of Wash Block Buffer was mixed with 12 μL of αRb-12x-SunTag-GFP and 12 μL of αMs-12x-MoonTag-GFP and incubated at room temperature for 30 minutes. The supernatant was removed from the cells, and the Block SunTag/MoonTag mix was added to the cells for incubation at 4°C for 1 hour. If multiple antibody combinations are to be run in parallel, split the 120 μL of Block SunTag/MoonTag mix between these samples.

Samples were washed twice in 100 μL of Wash Block Buffer and resuspended in 1500 μL of 300-Wash Block Buffer. Here, if there are multiple antibody combinations, bring each sample up in a volume of 300-Wash Block Buffer proportional to the fraction of that plate dedicated to that sample. Next, 15 μL of sample was added to each well of a 96-well plate (Eppendorf, cat. no. 0030129504) with frequent mixing in between additions to prevent bead bound cells from settling to the bottom of the tube. 1.5 μL from a differentially barcoded αSun-Tn5 PAGE plate was added to each well with a multi-channel pipette, and the plate was sealed and incubated at 4°C for 1 hour.

The plate of cells was spun down for one second at 1,200 RPM and placed on a 96-ring low elution magnet (ALPAQU cat. no. A000350). The wells were washed two times with 100 μL of 300-Wash Block Buffer and brought up in 50 μL of Tagmentation buffer. The plate was sealed and placed on a thermocycler, and tagmentation was performed at 37°C for 1 hour.

The plate was then placed back on the magnet stand, washed twice with 100 μL of 300-Wash Block Buffer and brought up in 15 μL of wash block buffer. 1.5 μL of differentially barcoded αMoon-Tn5 PAGE plate was added to each well with a multi-channel pipette and the plate was sealed and incubated at 4°C for 1 hour.

The plate of cells was spun down for one second at 1,200 RPM and placed back on the 96-well magnet stand. The wells were washed two times with 100 μL of 300-Wash Block Buffer and brought up in 50 μL of Tagmentation buffer. The plate was sealed and placed on a thermocycler, and tagmentation was performed at 37°C for 1 hour.

Tagmentation Buffer was removed from each well on the magnet stand, the plate was transferred to a non-magnetic rack, and all wells were washed and combined using 8 aliquots of 100 μL of Wash Buffer via multichannel pipetting each row together. Another round of 8 x100 μL was used to clean up the rows and the total 1.6 mL was combined and placed in a tube on a magnet stand. The pool was washed once with 500 μL of Wash Buffer and resuspended in 500 μL of Wash Buffer.

For size filtering, the 500 μL of Wash Buffer was passed through a 20μm filter (pluriSelect Cat no. 43-10020-40) at 100 × *g* for 3 minutes. DAPI staining was performed by adding 10 μL of 10,000 × DAPI diluted 2:198 in TAPS Buffer. Nuclei were counted on a Countess slide. Based on the cell concentration, cells were diluted to 10 cells per 35 nL in 10 mM TAPS, and 2:198 DAPI was added to a final concentration of 1x. Once in TAPS buffer cells will start to lyse and clump, and so it is important to start the dispense on the ICELL8 quickly. 80 μL were added into wells A1-D2 on the 384 source plate and 35 nL were dispensed into each well of a ICELL8 350v chip. The chip was then dried with a paper wipe, sealed, spun for 1 minute at 1,200 × *g* and imaged. 0.19% SDS Release Buffer was prepared (616 μL 10 mM TAPS, 152 μL 1% SDS, 32 μL Thermolabile Proteinase K). 35 nL of 0.19% SDS + thermolabile proteinase K in TAPS was dispensed on the chip, which was dried, sealed, and spun in the same manner. The chip was then incubated at at 37°C for 1 hour and then 58°C for 1 hour. The chip was spun again, and 35 nL of 2.5% Triton X-100 was dispensed, and the chip was dried, sealed, and spun in the same manner. 35 nL i5 primer was dispensed, and the chip was dried, sealed, and spun in the same manner. 35 nL i7 primer was dispensed, and the chip was dried, sealed, and spun in the same manner. KAPA PCR Mix was prepared with 450 μL 5 X HiFi buffer, 247.5 μL ddH_2_O, 67.5 μL 10 mM dNTP, 45 μL KAPA HiFi polymerase. 100 nL of KAPA PCR Mix was dispensed on the chip, and the chip was dried, sealed, and spun in the same manner. The chip was then placed into the custom thermocycler, and PCR was performed using the following thermal profile: 58°C for 5 min (gap fill), 72°C for 10 min, 98°C for 45 seconds, followed by 13 cycles of (98°C for 15 seconds, 60°C for 10 seconds, 72°C for 5 seconds), ending with 72°C for 1 minute and hold at 12°C. After amplification the chip was spun in a collection kit for 2 minutes at 1,200 RPM. The DNA pool was then put through an Ampure bead cleanup with 1.3 X size selection and eluted in 100 μL Tris-HCl pH 8.0. This was followed by an Ampure bead cleanup with 0.9 X size selection to remove the PCR product of “internal tagmentation” that results from one adapter tagmenting into another. DNA was eluted in 25 μL Tris-HCl pH 8. Final libraries were analyzed on a TapeStation. Sequencing is performed on an Element Aviti or Illumina Platform using the appropriate custom primers.

### Standard CUT&Tag

CUT&Tag was performed using a direct-to-PCR protocol like the one previously described^27^. Briefly, whole cells were spun down at 600 × *g* and brought up in 500 µL Wash Buffer containing Triton-X 100 for permeabilization (48 mL ddH_2_O, 1 mL 1M HEPES pH 7.5 [20 mM], 1.5 mL 5M NaCl [150 mM], 12.5 µL 2M Spermidine [0.5 mM], 250 µL of 10% Triton-X-100 [0.05%],1 Roche Complete EDTA-free tablet). ConA magnetic beads (Bangs, cat. no. BP531) were prepared by washing 5 µL per sample, 2 times with 500 µL Binding Buffer (400 µL of 1 M HEPES-KOH pH 7.9 [20 mM], 200 µL of 1 M KCl [10 mM] 20 µL of 1 M CaCl_2_ [1 mM], and 20 µL of 1 M MnCl_2_ [1 mM], bring up to 20 mL with nanopure water). ConA Beads were resuspended in 100 µL of Binding Buffer and were added to the whole cell suspensions in 3-33 µL increments, the sample was mixed by inversion after each addition. Bead binding was allowed to proceed for 10 min at room temperature, and the supernatant was removed on a magnet stand. Then cells were washed once with 500 µL of Wash Buffer and brought up in 15 µL of Antibody Buffer per CUT&Tag reaction/replicate. 1.5 µL of the rabbit or mouse primary antibody (e.g, rabbit-anti H3K27me3; Cell Signaling Technologies Cat No. 9733S). Antibody binding was allowed to proceed overnight at 4°C. The next day, cells were washed twice with 200 µL of Wash Buffer and brought up in 25 µL of Wash Buffer and 1.5 µL of secondary antibody (either guinea pig anti-rabbit IgG antibody; Antibodies Online Cat No. ABIN101961 or rabbit anti-mouse IgG antibody; Abcam Cat No. Ab46540) is added to each tube and incubation is allowed to proceed at 4°C for one hour. Cells were then washed twice with 200 µL of Wash Buffer and brought up in 15 µL of 300-Wash Buffer (48 mL ddH_2_O, 1 mL 1M HEPES pH 7.5 [20 mM], 3 mL 5M NaCl [300 mM], 12.5 µL 2M Spermidine [0.5 mM], 250 µL of 10% Triton-X-100 [0.05%], 1 Roche Complete EDTA-free tablet) and 1.5 µL of pA-Tn5 purified and loaded according to standard protocols (dx.doi.org/10.17504/protocols.io.8yrhxv6) was added and incubation is allowed to proceed at 4°C for one hour. Cells were then washed twice with 200 µL of Wash Buffer and brought up in 50 µL of tagmentation buffer (1 mL 300-Wash Buffer + 10 µL 1M MgCl_2_ [10 mM]), and tagmentation was carried out at 37°C for 1 hour on a thermocycler with a heated lid. Cells were then washed one time on the magnet stand with 200 µL Wash Buffer and one time with TAPS Buffer (500 µL 1M TAPS pH 8.5 [10 mM] in 49.5 mL ddH_2_O), and then brought up in 5 µL SDS Release Buffer (220 µL 10 mM TAPS, 25 µL 1% SDS [0.1% final], 5 µL Thermolabile Proteinase K). Samples are incubated on the thermocycler with a heated lid at 37°C for 1 hour and 58 degrees for one hour. The SDS is then quenched by addition of 4 μL of 1.5% Triton-X-100 (75 µL of 10% TritonX-100 solution in 425 µL of nanopure H_2_0). Each sample is then diluted by addition of 24 µL of nanopore H20, followed by 18 µL of PCR “Master Mix” (for each sample: 10 µL 5 X HiFi buffer, 5.5 µL ddH_2_O, 1.5 µL 10 mM dNTP, 1 µL KAPA HiFi Polymerase). Then add 2 µL of a barcoded Forward primer (10 µM) and 2 µL of a barcoded Reverse primer (10 µM) to each sample, ensuring that each sample has at least one distinct barcode. Mix by vortexing. Then place samples in Thermocylcer and run the following protocol with a heated lid at 105°C: 58°C for 5 min (gap fill), 72°C for 5 min, 98°C for 45 seconds, followed by 14 cycles of (98°C for 15 seconds, 60°C for 10 seconds, 72°C for 1 seconds), ending with 72°C for 1 minute and hold at 12°C. Samples are then purified using a standard Ampure bead cleanup with a 1.3 X size selection and then run on a Tapestation and pooled at equimolar concentrations for sequencing on the Element Aviti Platform.

### Single-cell CUT&Tag

To enable direct comparisons to sciCUT&Tag data generated in the same experiment, we dedicated 8 wells of the 96-well plate used for CoCUT&Tag to samples bound with a single antibody, and the appropriate guinea pig anti-rabbit or rabbit anti-mouse antibody secondary antibody. Samples were then distributed into the 96 well plate in parallel to the CoCUT&Tag samples, in 15 µL of 300-Wash Buffer per well. Cells were then bound by 1.5 µL of differentially barcoded pA-Tn5 prepared as previously described^45^, ensuring the pA-Tn5s were loaded with barcoded oligos that were compatible with those used for the CoCUT&Tag samples. After the first round of tagmentation, the standard sciCUT&Tag samples were removed from the plate and pooled, and these cells were then added back to the pool of CoCUT&Tag samples after they had completed the second round of tagmentation. This combination of sciCUT&Tag and CoCUT&Tag cells was then distributed on the ICELL8 nanowell dispenser together for PCR indexing.

### Sequencing

All libraries used in this study were sequenced on the Element Aviti platform. Standard CUT&Tag libraries are compatible with the standard primer mix supplied as part of the Element Aviti flow cell kit. Sequencing the CoCUT&Tag libraries on the Element Aviti platform requires the custom primers: Read1_ seq_primer (GAGGACGGCAGATGTGTATAAGAGACAG), R e a d 2 _ s e q _ p r i m e r (GTCTCCGCCTCAGATGTGTATAAGAGACAG), Index1_ seq_primer (CATCTGAGGCGGAGACGGTG), Index2_ seq_primer (AATGATACGGCGACCACCGAGATCTACAC). The read corresponding to Index1 must be 43 cycles, the read corresponding to Index2 must be 37 cycles, and the rest of the cycles were split evenly between genomic read1 and genomic read2.

### Preprocessing of CoCUT&Tag Sequencing

The preprocessing of CoCUT&Tag sequencing involves demultiplexing, rewriting cell barcodes, trimming adapters, alignment, bed file generation, and creating coverage-normalized bigwig files.

In our combinatory index scheme, cell barcodes are determined by the combination of Tn5 adaptor barcodes and iCell8 well index barcodes. The set of barcoded Tn5 adapters PAGE-s5-1, PAGE-s7-1 were used to load anti-SunTag-Tn5, and the set of barcoded Tn5 adapters PAGE-s5-2 and PAGE-s7-2 were used to load anti-MoonTag-Tn5 and both sets were introduced during the Tn5 tagmentation steps in a 96 well plate. The 72-by-72 iCell8 well index barcodes (p5 and p7) were subsequently added by PCR in the iCell8 nanowell dispenser. Each cell contained a single combination of iCell8 p5 and p7 barcodes, but could be associated with four possible combinations of Tn5 adaptor barcodes: PAGE-s5-1 + PAGE-s7-1 reads are specific to sites bound by the rabbit-IgG-SunTag-labeled protein, PAGE-s5-2 + PAGE-s7-2 reads are specific to sites bound by the mouse-IgG-MoonTag-labeled protein, PAGE-s5-1 + PAGE-s7-2, and PAGE-s5-2 + PAGE-s7-1 reads are specific to the co-occupancy sites where the proteins targeted by the rabbit and mouse antibodies are bound together. Those combinations of Tn5 adaptor barcodes were used for demultiplexing the FASTQ file using the sciCTextract program (https://github.com/mfitzgib/sciCTextract) to generate mono-occupancy and co-occupancy FASTQ files. Within each FASTQ file, cell barcodes, consisting of the combined iCell8 well index barcodes and Tn5 adaptor barcodes, were stored in the read header. To simplify the matching of reads and cells across modalities in downstream analyses, we revised the cell barcode format by replacing the four Tn5 adaptor barcode combinations with a well ID from the 96-well plate used in the Tn5 tagmentation step. In this format, reads originating from the same cell have the same barcode and are split across four fastq files, (1) reads specific to the rabbit-IgG-SunTag-labeled protein, (2) reads specific to the mouse-IgG-MoonTag-labeled protein, (3) a first set of co-occupancy reads and (4) a second set of co-occupancy reads. By splitting the co-occupancy reads across two fastq files, we preserve information regarding which paired end of the read corresponds to each target protein, but for most applications the co-occupancy reads can be combined for downstream analysis.

After demultiplexing and barcode rewriting, adapter trimming was performed with Cutadapt (v4.3) using paired-end mode, a minimum read length of 20 bases, and the adapter sequence “CTGTCTCTTATACACATCT” applied to both read 1 and read 2. Trimmed read pairs were aligned using Bowtie2 (v2.5.4) to the hg38 indexed reference genome. The alignment command used the options --very-sensitive-local, --soft-clipped-unmapped-tlen, --no-mixed, --no-discordant, --dovetail, --phred33, -I 10, and -X 1000. Aligned SAM files were converted to BAM files with Samtools (v1.21). The BAM files were then converted to paired-end BED fragments with the Bedtools (v2.30.0) “bamtobed -bedpe” command. Fragment records were reformatted to retain genomic coordinates and barcode-derived identifiers and were then sorted and written as compressed BED files. The single-cell CoCUT&Tag data was collapsed after sorting to remove duplicate reads because the likelihood of obtaining biological duplicates from the same cell is exceedingly low. Duplicate reads were retained in the bulk CoCUT&Tag data. Using the BED files, we generated normalized coverage tracks. Total fragment coverage was first calculated as the summed fragment length across all BED intervals. A per-sample scaling factor was then computed as (1 /Total Coverage) x 10^10. Bedtools genomecov was used with the -bg option to generate coverage-normalized BedGraph files using the per-sample scaling factor together with a chromosome sizes file corresponding to the hg38 reference genome. UCSC bedGraphToBigWig (v2.10) was used to convert the BedGraph output to BigWig format.

We developed a Nextflow pipeline to conduct the preprocessing of single-cell CoCUT&Tag available at: https://github.com/vari-bbc/sciCT_pipeline.

### H3K27ac normalized CoCUT&Tag tornado plots and average plots

To enable quantitative comparisons of the co-occupancy reads between our H3K4me3-H3K27ac and H3K27me3-H3K27ac paired CoCUT&Tag samples, we first concatenated and sorted the bed files from the mono-occupancy and co-occupancy reads from two biological replicates. We then calculated a single scaling factor for each sample computed as (1/Total H3K27ac Coverage) x 10^10. This scaling factor was then applied during the Bedtools genomecov generation of BedGraphs for the mono-occupancy and co-occupancy files corresponding for the paired samples. UCSC bedGraphToBigWig (v2.10) was used to convert the BedGraph outputs to BigWig format. We then called peaks on the H3K27me3, and H3K4me3 and H3K27ac bedfiles using SEACR^47^ using the settings “non” normalized with an FDR threshold of 0.01 and “stringent”. H3K4me3-H3K27ac and H3K27me3-H3K27ac shared peaks were identified by merging the two peak files and then using the bedtools intersect -u command to keep only the merged peaks that overlapped with peaks in each of the original peak sets. The H3K4me3 and H3K27ac specific peak sets and the H3K27me3 and H3K27ac specific peak sets were then generated by using the bedtools intersect -v command to remove any peaks that overlapped with the corresponding shared peak set. We then used deeptools version 3.5.2 to generate a matrix file from the H3K27ac-scaled bigwigs using the parameters computeMatrix reference-point -R shared_ peaks.bed specific_peaks_1.bed specific _peaks_2.bed -S monooccupancy_1.bigwig monooccupancy_2.bigwig cooccupancy.bigwig -a 5000 -b 5000 -bs 50 --sortRegions keep --missingDataAsZero --referencePoint center. The matrix files were then used as input to generate the corresponding tornado plots using the plotHeatmap function and the corresponding average plots using the plotProfile function.

### Single-cell CoCUT&Tag quality control and doublet removal

Single-cell chromatin profiles were processed in R using ArchR^22^ (v1.0.3) with the hg38 reference genome. Arrow files were generated from compressed BED files using createArrowFiles(). For the mono-occupancy datasets, Arrow file generation was performed with minTSS = 0, minFrags = 100, maxFrags = Inf, nChunk = 1, and excludeChr = “chrM”. For co-occupancy datasets, the same parameters were used except that minFrags = 0, allowing all fragments to be retained at the Arrow-generation stage and to be filtered later using the barcodes from cells that pass the mono-occupancy quality control. Each cell has three modalities, they are two mono-occupancy modalities and one co-occupancy modality as we do not differentiate between the two heterogeneous Tn5 adaptor barcode combinations in the co-occupancy case. If a cell has more than 250 fragments in one of the three modalities, it was retained for downstream analysis.

Using corresponding arrow files, we created ArchR projects for each modality. For each project, a TileMatrix was generated after excluding reads from chrM, chrX, and chrY. Different tile sizes were used for different assay classes: H3K27me3 mono-occupancy projects were tiled at 50-kb resolution, whereas H3K4me1, H3K4me2, and H3K4me3 mono-occupancy projects were tiled at 5-kb resolution. Co-occupancy projects were also processed with 50-kb tiles. Low-dimensional representations were then computed from these tiled matrices using LSI (latent semantic indexing) with addIterativeLSI(), followed by Harmony correction with addHarmony() and UMAP embedding addUMAP() using default parameters.

Single-cell profiles of peripheral blood mononuclear cells (PBMCs) were generated from a single donor, and putative doublets were identified used the ArchR function addDoubletScores(). This approach generates synthetic doublets and projects them with observed cells into a low-dimensional space using latent semantic indexing (LSI; dimensions 1–15), allowing each cell to be assigned a doublet enrichment score based on its proximity to simulated doublets. Because the PBMC experiments compared multiple versions of the SunTag and MoonTag scaffolds, as well as CoCUT&Tag and sciCUT&Tag reactions performed with different antibodies, the number of cells obtained for any individual condition was limited. To increase cell numbers and improve doublet scoring, Arrow files were generated from concatenated, sorted, and compressed BED files corresponding to all data generated with a given antibody.

Thus, standard sciCUT&Tag data and data generated with 4x, 12x, or 24x SunTag or MoonTag scaffolds were combined prior to doublet scoring. As an initial quality control step, H3K4me3 clusters with unusually low TSS scores were removed. Next, addDoubletScores() was applied separately to the combined H3K27me3 and H3K4me3 Arrow files. To remove doublet-enriched populations, cells were first clustered in ArchR and the fraction of cells with elevated doublet enrichment scores was calculated for each cluster. For each histone mark a DoubletEnrichment score was defined, and cells with scores above this threshold were classified as putative doublets. The percentage of putative doublets was then calculated for each cluster. Clusters were rank-ordered by doublet percentage, and clusters in which the fraction of putative doublets exceeded a defined threshold were removed from the ArchR project prior to downstream analysis.

For experiments on human bone marrow, we mixed an equal number of cells from four different donors, and used SouporCell^20^ (v2.5) to generate genotype-based doublet calls for all of the barcodes included in one or more of the mono-occupancy or co-occupancy ArchR projects. For this, aligned BAM files were updated by adding cell barcode tags CB, extracted directly from the read names. Then, the BAM files were deduplicated using samtools (v1.21) sort, fixmate, and markdup, with duplicate marking performed in a cell-barcode-aware manner via --barcode-tag CB to avoid collapsing reads across different cells. The deduplicated BAM files were merged, coordinate-sorted, and indexed with samtools. The merged BAM, a barcode whitelist file including all cells from any of the ArchR projects, and the same reference genome FASTA file used during alignment were then supplied to souporcell_pipeline.py of SouporCell to do the single-cell genotype clustering. Souporcell was run in no-UMI mode with remapping disabled, using parameters: --no_umi True, --skip_remap True, and --ignore True, together with a user-specified genotype cluster count (-k) determined by the mixed donor number. The output of Souporcell was then applied to the ArchR project to identify doublet-enriched neighborhoods. Cells were partitioned into 100 groups in Harmony space using k-means clustering, and the fraction of cells with souporcell status of “doublet” was calculated for each group. K-means groups with more than 15% souporcell-labeled doublets were considered doublet-enriched and all cells in those groups were removed from all three modalities. In downstream QC summaries, the combined fraction of souporcell doublets and unassigned cells was also inspected at the cell type cluster level to detect residual problematic populations. To run Souporcell, we adapted the nextflow pipeline associated with our previous sciCUT&Tag publication (https://github.com/vari-bbc/souper-star)^45^.

### Peripheral blood mononuclear cell annotation and embedding fidelity metrics

After doublet removal, gene score matrices and a reads in genomic bins matrix was extracted ArchR projects corresponding to H3K4me3 and H3K27me3 along with cell metadata and low-dimensional embeddings. These data were then imported into Python and processed using Scanpy (v1.x) and associated libraries for multimodal AnnData integration.

Weighted nearest neighbor (WNN) embeddings were generated using a custom integration pipeline. Briefly, modality-specific dimensionality reduction was first performed independently for each mark (principal component analysis for gene score matrices and latent semantic indexing for tile-based profiles). K-nearest neighbor graphs were then constructed for each modality using scanpy.pp.neighbors(). Cell-specific modality weights were learned by comparing the consistency of each cell’s local neighborhood across modalities, allowing adaptive weighting of H3K4me3 and H3K27me3 contributions. These modality-specific graphs were then combined into a unified WNN graph, which was used to compute a joint embedding using scanpy.tl.umap() and to perform clustering using the Leiden algorithm (scanpy. tl.leiden()).

The resulting clusters were used as the primary cell-state definitions for downstream analysis. Cluster identities were assigned based on marker genes identified from the H3K4me3 GeneScoreMatrix in ArchR using getMarkerFeatures() with Wilcoxon testing (FDR ≤ 0.01, log2FC ≥ 1.25), including bias correction for TSSEnrichment and log10(nFrags). For each cluster, the top-ranking H3K4me3 marker genes were used to annotate canonical PBMC populations based on enrichment of known cell-type specific marker genes.

To evaluate the fidelity of H3K4me3 and H3K27me3-based embeddings, we compared the proximity of matched profiles of the same histone mark profiled twice within the same cells. For H3K4me3, matched profiles were generated using the rabbit CST 9751 antibody and mouse Diagenode C15200152 antibody, and for H3K27me3, matched profiles were generated using the rabbit CST 9733 antibody and the mouse and BPS Bioscience 25243 antibody. For each matched pair, we calculated the Euclidean distance between the two profiles in Harmony space using dimensions 1-15. To place this distance in the context of the local embedding density, we then sampled up to 5,000 other cells from the same two-sample comparison and calculated the fraction of sampled cells that were closer to each cell than its true matched partner. This value averaged to generate a pair-closeness score, S_pair, where lower values indicate that matched profiles occupy unusually close positions in the embedding relative to unrelated cells. The distributions of S_pair values were then compared for H3K4me3-H3K4me3 and H3K27me3-H3K27me3 matched profiles using box plots and empirical cumulative distribution, providing a quantitative measure of embedding fidelity for each histone mark.

### Annotation of human bone marrow CoCUT&Tag

After removal of putative doublets, dimensionality reduction and clustering were repeated with default parameters in ArchR. Cluster quality was evaluated in multiple ways. First, cluster correspondence across modalities was assessed by comparing cluster consistency. Second, a GeneScoreMatrix was used to identify cluster-enriched marker genes for each histone modification, and the top upregulated marker genes from each cluster were compared with an external bone marrow scRNA-seq dataset annotated with the surface antigen information to determine cell type identity^21^. Third, cluster-level quality metrics, including fragment-size distributions, transcription start site enrichment (TSSEnrichment), number of fragments (nFrags), donor representation, antibody, and residual doublet or unassigned fractions, were reviewed to identify unstable or artifactual populations. Clusters judged to be inconsistent across marks or enriched for odd cluster-level quality metrics comparing to majority clusters were removed from the relevant ArchR projects before the next round of clustering. H3K27me3 cluster labels were propagated to matched cells in the other modalities as a shared cell-type annotation. A second round of marker-gene calling and scRNA-seq-reference mapping was then performed on the refined ArchR projects.

For clustering consistency between modalities, we compared the clustering result of the two mono-occupancy modalities for each cell and created a cluster-cluster overlap matrix. Using this overlap matrix, we visually assessed the clustering consistency between the two modalities with a heatmap.

For comparing the CoCUT&Tag cluster assignments to scRNA-seq cell-type annotations, the cluster-associated marker genes were identified from the H3K4me3 GeneScoreMatrix with getMarkerFeatures(), with Wilcoxon testing (FDR <= 0.01 and Log2FC >= 1.25), including bias correction for TSSEnrichment and log10 nFrags. For each CoCUT&Tag cluster, we selected the top 50 genes with the most significant FDR values as the marker gene set, then determined their expression in each cell cluster of the reference scRNA-seq using the normalized RNA expression without scaling. After applying z-score scaling and averaging RNA expression, we created a matrix to compare CoCUT&Tag clusters with scRNA-seq-annotated cell types. Each entry in the matrix represents the average scaled mean expression of marker genes for each CoCUT&Tag cluster across the specific scRNA-seq-annotated cell types. The matrix was visualized with a heatmap, and we conducted the same comparison for H3K4me1 and H3K4me2.

### Gene-score imputation

To get smoothed imputed gene scores, the original gene-score matrices were extracted from each ArchR project as gene-by-cell matrices. Gene-score imputation was then performed using ArchR imputation weights derived from the H3K27me3 Harmony embedding, yielding a smoothed H3K27me3 gene-score matrix together with cross-mark imputed matrices for mono-occupancy H3K4me1, H3K4me2, H3K4me3 and co-occupancy H3K27me3-H3K4me1, H3K27me3-H3K4me2, and H3K27me3-H3K4me3 combinations. These imputed gene scores were used for gene score related analysis.

### Pseudo-bulk generation

After obtaining the cell annotations, we created a pseudo-bulk gene score matrix by averaging the gene scores for each cell-type cluster per gene. We also generated pseudo-bulk bed files and cell-number-normalized bigwig files for each cell type. Firstly, we create a bed file for each histone modification mono-occupancy or co-occupancy per cell-type cluster. Then a per cell-type cluster scaling factor was computed as (1 /Total cell number per cluster) x 10000. Bedtools genomecov was used with the -bg option to generate scaled BedGraph coverage using a chromosome sizes file derived from hg38 reference genome, and UCSC bedGraphToBigWig was used to convert the BedGraph output to BigWig format.

### Bone marrow CoCUT&Tag chromatin-state assignment

Chromatin-state assignment was conducted on cell-type level pseudo-bulk gene scores to classify each gene per cell state as active, repressive, bivalent, or unmarked. To guide threshold selection, we modeled the distribution of pseudo-bulk gene scores for each mark using a two-component Gaussian mixture model on log10(gene score + 0.1) with the mclust() in R package mclust (v6.1.2). This provided the mean, standard deviation, and mixture weight of each component, defining two weighted component densities. The threshold was set at the intersection of these densities, representing the point of equal likelihood between low- and high-signal components. These thresholds were used for subsequent classification. Chromatin state was assigned separately for each gene in each cell-type cluster for each H3K4 methylation state paired with H3K27me3 (i.e., H3K27me3-H3K4me1, H3K27me3-H3K4me2, and H3K27me3-H3K4me3). Genes were classified as bivalent if the H3K4 signal, H3K27me3 signal, and co-occupancy signal all exceeded the threshold; as active if the H3K4 signal was above the threshold and H3K27me3 was below; as repressive if H3K27me3 was above the threshold and the H3K4 signal was below; as unmarked if both signals were below the threshold; or as other if genes did not fall into any of these categories. Genes classified as “other” were excluded from further analysis. To derive a multi-mark consensus, categorical assignments from the H3K27me3-H3K4me1, H3K27me3-H3K4me2, and H3K27me3-H3K4me3 marker pairs were compared for each gene and cell type. A consensus state was defined as concordant assignments in at least two of the three marker pairs in a given cell type.

### Bone marrow trajectory analysis

We defined the root for CoCUT&Tag trajectory analysis by re-clustering cells in the HSC/MPP cluster and the connected lymphoid, myeloid, and erythroid cell clusters. We performed dimension reduction with addIterativeLSI(), batch effect correction with addHarmony(), and clustering on Harmony embeddings using addClusters() with bias filtering enabled and customized parameters: filterBias = T, biasCluster = 0.2, baisCol = ‘nFrags’, baisEnrich = 5, biasProportion = 0.10, bias Pval = 0.05. We then propagated the cluster labels assigned to the H3K27me3 modality to the matched cells in the H3K4me1, H3K4me2, and H3K4me3 embeddings. Using known stem-cell marker genes (*CD34, MYCN, MECOM, HOXB5, PRDM16*) and granulocyte-myeloid progenitor marker genes (*MPO* and *IRF*8), we identified an HSC/MPP subcluster with more stem cell-like features, and another subcluster that was biased towards the myeloid lineage.

The new stem cell-like subcluster was assigned as the root and the myeloid biased HSC/MPP subcluster was assigned as the start of the myeloid lineage. We inferred trajectories for the B-cell, myeloid and erythroid lineages using the ArchR addTrajectory() function with default parameters. Pseudotime was determined by ranking the cells along the trajectory and scaling it from 0 to 100. For the gene module CoCUT&Tag pseudotime gene score scatter plot, the gene scores were first standardized by calculating the z-score for each gene across cells. The median z-score of the gene list for each cell was then used as the gene module score. In the CoCUT&Tag pseudotime gene score scatter plot, the imputed gene score was used, and a smooth line was fitted using LOESS model with a span of 0.3.

In each lineage we determined gene activation timing by identifying the earliest pseudotime position (from 0 to 100) where a gene’s scaled, smoothed activity reaches at least 50% of its peak in a lineage. Specifically, for each gene, we smoothed the imputed gene score using the loess model with the default span value via the R loess() function by the pseudotime, using the predicted values. Non-positive values are set to 0. We then rescaled each gene’s activity with z-score to a range between 0 and 1 by subtracting the minimum and dividing by the range. The first pseudotime index where the scaled value exceeds 0.5 is defined as the activation time.

For gene expression pseudotime analysis using the scRNA-seq dataset^21^, we utilized the trajectory information from the original Seurat object’s metadata and assigned pseudotime based on the scaled rank (from 1 to 100) of cells within each trajectory. For the scRNA-seq pseudotime gene expression scatter plot, gene expression values were used from the data slot, which was gene count divided by cell’s total counts, multiplied by the scale factor, and then transformed as log1p. In the scatter plot, the smooth line was fitted using LOESS model with a span of 0.15. For the unscaled gene module pseudotime expression scatter plot, the average expression of a gene module was calculated per cell. In the scaled case, this average expression was standardized by subtracting the mean and dividing by the standard deviation across cells. And in the gene module pseudotime expression scatter plot, the smooth line was fitted using LOESS model with a span of 0.7.

### Classification of cis-regulatory elements

First we defined gene domains from GENCODE v38 annotations by (1) filtering for protein-coding transcripts, (2) collapsing multiple transcripts into a single gene interval by selecting the most upstream TSSs and most downstream transcription end sites (TESs), and (3) scanning chromosome by chromosome to identify the nearest upstream and downstream non-overlapping neighboring gene intervals for each gene. The upstream boundary of each gene domain was set to the end of the nearest upstream non-overlapping gene, and the downstream boundary was set to the start of the nearest downstream non-overlapping gene; genes at chromosome edges were assigned boundaries based on the minimum or maximum annotated coordinate on that chromosome. Invalid intervals were removed, yielding a final BED-like gene-domain annotation containing chromosome, domain start, domain end, and gene name for downstream assignment of regulatory elements to genes.

To identify candidate cis-regulatory elements, peaks were called on the cluster specific H3K4me2 bed files from the HSC/MPPs, and the B, myeloid and erythroid lineages. Then, for each lineage, all summit bedfiles were loaded, concatenated, and sorted by chromosome and genomic coordinate. Summit calls occurring within 100 bp of one another on the same chromosome were collapsed using a greedy sweep, retaining the summit with the highest MACS3 score. Each retained summit was then converted into a fixed-width 200 bp BED interval centered on the summit.

We restricted our initial analysis to elements associated with lineage-specific activation. The previously defined gene-domain annotation was filtered to include the unmarked-to-active, and bivalent-to-active gene sets for each lineage. These gene domains were used to select the 200 bp summit windows using BedTool.intersect(u=True). Retained B, erythroid, and myeloid peak sets were then concatenated and merged with BedTool.merge() to generate a non-redundant cross-lineage peak set. Some genes are activated in multiple lineages, and the H3K4me2 peak summits may be different in each lineage. To account for this, merged intervals greater than 200 bp were reassigned to a single representative summit by intersecting them back to the complete set of original MACS3 summits and selecting the overlapping summit with the highest MACS3 score. These representative summits were then re-centered into fixed 200 bp windows, yielding a final standardized summit-centered feature set for downstream motif analysis. Read counts for each chromatin modality, including H3K27me3, H3K4me2, H3K27me3/H3K4me2 co-occupied bivalent reads, H3K4me3, and H3K4me1 were then quantified over the final combined 200 bp summit-centered feature set using BedTool.intersect(c=True), which reports the number of overlapping reads per interval. Total reads were for each peak were normalized by the total number of reads in the cluster and scaled to counts per million reads. The resulting values were assembled into a feature matrix in which rows corresponded to the standardized 200 bp summit-centered intervals and columns corresponded to cluster-histone-mark combinations (e.g., HSC-H3K27me3, HSC-H3K4me2, or HSC-Biv). Genomic coordinates and peak identifiers were retained with the matrix for downstream grouping and motif analysis.

Low- and high-signal outlier features were removed by calculating the cumulative signal for each peak as the sum across all cluster-histone-mark columns. Peaks in the lowest 5% and highest 5% of the cumulative signal distribution were removed. The retained features were clustered based on their chromatin signal profiles across cell states and histone marks. Signal columns were standardized column-wise using StandardScaler, without an additional log transformation. To prevent marks with larger aggregate signal from dominating the analysis, columns were grouped by histone mark, and each mark-specific block was rescaled to have the same overall matrix norm, using the median block norm as the target. Principal component analysis was then performed with sklearn.decomposition.PCA, and the first 16 principal components were used for graph construction. A k-nearest-neighbor graph was generated with NearestNeighbors using 30 neighbors, with edge weights defined as the inverse Euclidean distance between features. Leiden clustering was then performed on this weighted graph using leidenalg with an RBConfigurationVertexPartition and resolution parameter of 4. Finally, the same PCA representation was embedded in two dimensions using UMAP with 40 neighbors, a minimum distance of 0.25, and Euclidean distance, and feature clusters were visualized in UMAP space.

Lineage-restricted enhancer classes were further refined by a second round of clustering. After the initial global clustering of summit-centered features, promoter-associated clusters and lineage-specific enhancer-associated clusters were selected, and principal component analysis was repeated using sklearn.decomposition.PCA, with up to 15 principal components retained depending on the number of available peaks and dimensions. A new weighted k-nearest-neighbor graph was constructed from the subset PCA space using 30 neighbors, and Leiden clustering was repeated with a lower resolution parameter of 0.5 to identify subgroups within each class of elements. The reclustered features were visualized using UMAP with 40 neighbors, a minimum distance of 0.25, and Euclidean distance. The resulting clusters were then assigned biologically interpretable labels that were used to overwrite the original global cluster annotations. This produced the final cis-regulatory element annotations while preserving the original feature matrix.

Promoter overlap was quantified for each final feature cluster using a TSS-centered promoter annotation. A BED file of hg38 transcription start sites was expanded by 2 kb on each side using pybedtools.BedTool.slop(), with the hg38 genome index used to constrain intervals to chromosome boundaries, and the resulting promoter windows were sorted. The final peak feature set was then converted to a BedTool with peak identifiers retained in the fourth column, and promoter-overlapping peaks were identified using BedTool.intersect(u=True). For each feature cluster, we then calculated the promoter fraction, defined as the number of promoter-overlapping peaks, over the total number of peaks.

### Cis-regulatory element enrichment and co-occurrence

Element-class enrichment was calculated by intersecting the final peak annotations with lineage-specific gene-domain groups. First, gene lists from B, myeloid, and erythroid unmarked-to-active and bivalent-to-active gene categories were loaded and partitioned into mutually exclusive classes, including lineage-exclusive as well as pairwise shared B– myeloid, B–erythroid, and erythroid–myeloid groups. The resulting groups were checked to confirm that no gene was assigned to more than one class. For each gene class, the corresponding gene-domain intervals were extracted from the gene-domain file and converted to sorted pybedtools intervals. The various cis-regulatory element peak sets split by classification were then intersected with each gene-domain class using BedTool.intersect(u=True), and overlapping peaks were counted by their final element classification. These counts were used to construct an observed element-class by gene-domain-class matrix. Expected counts were calculated from the product of the total number of peaks in each element class and the genome-wide fraction of peaks overlapping each gene-domain class. Enrichment was reported as log2 observed/expected with a pseudocount of 0.5. To visualize relative gene-domain preferences for each element class, log2 enrichment values were converted to row-wise softmax weights so that the gene-domain-class contributions for each element type summed to one, and promoter-or enhancer-like element labels were then plotted as stacked bar charts.

Element co-occurrence within gene domains was then calculated from the gene-domain by-element counts matrix. For each gene, we then recorded whether at least one peak from each element class overlapped its regulatory domain, generating a binary gene-by-element matrix; gene-domain length and the total number of element classes per gene were retained as additional annotations. Pairwise co-occurrence between element classes was tested across genes using Fisher’s exact test on the binary presence/absence matrix. For each element pair, a 2 × 2 contingency table was constructed from genes containing both elements, only the first element, only the second element, or neither element, and enrichment was summarized as a pseudocount-stabilized log2 odds ratio. P values were adjusted across all element pairs using Benjamini–Hochberg correction, and significant co-occurrences were defined at FDR < 0.05. Significant positive relationships were visualized as a co-occurrence network, in which nodes represented element classes_41_and edges represented significant positive co-occurrence relationships. For network visualization, edges were filtered to retain positive associations, further simplified by minimum node degree and top-weighted backbone edges, and plotted with node color indicating lineage-associated element identity and edge width reflecting the magnitude of co-occurrence enrichment.

To quantify the number of specific element classes and H3K4me1 or H3K4me3 signal intensity within gene domains, enhancer classes were combined into a single category and promoter classes were likewise combined. Element counts were summarized across gene groups and visualized as box plots. To compare element counts between unmarked-to-active and bivalent-to-active gene groups within each lineage, overlap-count distributions were tested with the Mann-Whitney U test, followed by Benjamini-Hochberg correction.

We then quantified the chromatin signal intensity of overlapping elements within each gene-domain class. For a given histone mark, signal columns corresponding to hematopoietic cell states were selected from the previously generated feature matrix, and each peak was assigned its global maximum signal across all cell type clusters. For enhancer analyses, H3K4me1 signal was used as the representative enhancer-associated mark, and for promoter analyses, H3K4me3 signal was used as the representative promoter-associated mark. Signal distributions were summarized across reorganized gene-domain classes using box plots. Differences in signal between unmarked-to-active and bivalent-to-active gene groups were tested within each lineage using Mann-Whitney U tests, with Benjamini-Hochberg correction across comparisons.

### Motif Enrichment Analysis

For motif enrichment controls, we generated standardized promoter and distal background feature sets from the same H3K4me2 summit-centered peak universe but elements that overlapped the bivalent-to-active and unmarked-to-active gene domains were excluded. Lineage-specific 200 bp MACS3 summit windows from all lineages were first concatenated and merged to create a global peak universe. Merged intervals were recalibrated back to fixed 200 bp windows. Intervals >200 bp were intersected back to the complete set of original MACS3 summits, and the highest-scoring overlapping summit was selected as the representative center. The resulting fixed-width 200 bp feature universe was split into promoter and distal sets by intersecting with hg38 TSS annotations expanded ±2 kb using pybedtools(). Features overlapping a TSS window were assigned to the promoter background set, while non-overlapping features were assigned to the distal background set.

For motif enrichment analysis, a curated non-redundant PWM set was generated from JASPAR2022 human and mouse motif annotations using TFBSTools, JASPAR2022, and monaLisa. Position frequency matrices were first retrieved for human and mouse using getMatrixSet() with species identifiers 9606 and 10090, respectively, and converted to position weight matrices with toPWM(). Human and mouse PWMs were then combined and collapsed by JASPAR base motif ID, removing version-specific duplicates. When duplicate motifs were present, one representative was retained by prioritizing motifs matching hematopoietic transcription factor names, followed by human motifs. To further reduce redundancy among highly similar motifs, pairwise PWM similarity was calculated using Pearson-based PWMSimilarity(), and a greedy pruning strategy was applied to remove motifs with similarity ≥0.95 to a higher-ranked representative.

Motif enrichment was performed across all distal element clusters using the curated JASPAR2022 PWM set with the monaLisa framework. For each cluster the promoter or enhancer elements were defined as foreground elements (fg), and compared against the those of the corresponding standardized background promoter and distal feature universe (by). Both foreground and background intervals were filtered to retain only sequences composed of canonical A/C/G/T bases. DNA sequences for foreground and background intervals were extracted from the hg38 genome using BSgenome.Hsapiens.UCSC.hg38, using the genomic intervals defined by the imported BED files and converted to GRanges objects. Motif enrichment was then computed using monaLisa::calcBinnedMotifEnrR(), applying a Fisher’s exact test to compare motif occurrences between foreground and background elements. For each cluster, motif enrichment results were stored as SummarizedExperiment objects and exported as tab-delimited tables containing motif IDs, motif names, log2 enrichment values, and −log10-transformed adjusted P values. Analyses were performed independently for each cluster.

### Statistics

Gene set overrepresentation analysis (ORA) was performed using the enrichGO() function from the R package clusterProfiler (v4.14.6), with the org.Hs.eg.db (v3.20.0) database. Wilcoxon rank-sum statistics were used for all unpaired data. The heatmap was plotted with pheatmap (v1.0.12) and other data visualization was done with ggplot (4.0.0) in R.

